# A comparison of marker gene selection methods for single-cell RNA sequencing data

**DOI:** 10.1101/2022.05.09.490241

**Authors:** Jeffrey M. Pullin, Davis J. McCarthy

**Affiliations:** Bioinformatics and Cellular Genomics, St Vincent’s Institute of Medical Research, Fitzroy, Australia; School of Mathematics and Statistics, Faculty of Science, University of Melbourne, Parkville, Australia; Melbourne Integrative Genomics, Faculty of Science, University of Melbourne, Parkville, Australia

## Abstract

The development of single-cell RNA sequencing (scRNA-seq) has enabled scientists to catalogue and probe the transcriptional heterogeneity of individual cells in unprecedented detail. A common step in the analysis of scRNA-seq data is the selection of so-called marker genes, most commonly to enable annotation of the biological cell types present in the sample. In this paper we benchmarked 56 computational methods for selecting marker genes in scRNA-seq data. The performance of the methods was compared using 10 real scRNA-seq datasets and over 170 additional simulated datasets. Methods were compared on their ability to recover simulated and expert-annotated marker genes, the predictive performance and characteristics of the gene sets they select, their memory usage and speed and their implementation quality. In addition, various case studies were used to scrutinise the most commonly used methods, highlighting issues and inconsistencies. Overall, we present a comprehensive evaluation of methods for selecting marker genes in scRNA-seq data. Our results highlight the efficacy of simple methods, especially the Wilcoxon rank-sum test, Student’s t-test and logistic regression. All code used in the evaluation, including an extensible Snakemake pipeline, is available at: https://gitlab.svi.edu.au/biocellgen-public/mage_2020_marker-gene-benchmarking.

## Introduction

Single-cell RNA sequencing (scRNA-seq) has enabled the high-throughput measurement of gene expression in single cells, enabling the interrogation of cell-type-specific changes in gene expression and regulation. Recently, decreases in cost and advances in protocol efficacy have led to a rapid increase in the number of scRNA-seq datasets used in biological research^[1]^. Over the same period, there has been a corresponding increase in the number of methods available to analyse scRNA-seq data. As of June 2022, there were over 1,250 tools available to perform various steps of an scRNA-seq data analysis^[2,3]^.

A ubiquitous step in the analysis of scRNA-seq data is the selection, by a computational method, of so-called marker genes. These marker genes are a small (typically ≤ 20) subset of genes that have expression profiles able to distinguish the subpopulations of cells present in the data. Most commonly, marker genes are selected with respect to a specific computational clustering of the data and are used to annotate, and understand, the biological cell type of the defined clusters (Figure 1a). Annotation of the cell-type of clusters is critical both to guide the clustering and to interpret the results of downstream analyses performed with respect to the clustering, such as differential expression testing^[4]^ or single-cell eQTL mapping^[5]^. In addition to cluster annotation, some authors treat the marker genes as new biological discoveries in their own right^[6]^, as candidates for further perturbation or differential analyses^[7]^, or as evidence for the quality of a particular clustering^[8]^. As well as their direct use, marker genes are a component of computational approaches that aim to annotate clusters automatically ^[9–11]^. Ultimately, marker genes provide a ‘human interpretable’ summary of the transcriptomic profiles of cell subpopulations.

**Figure 1:**
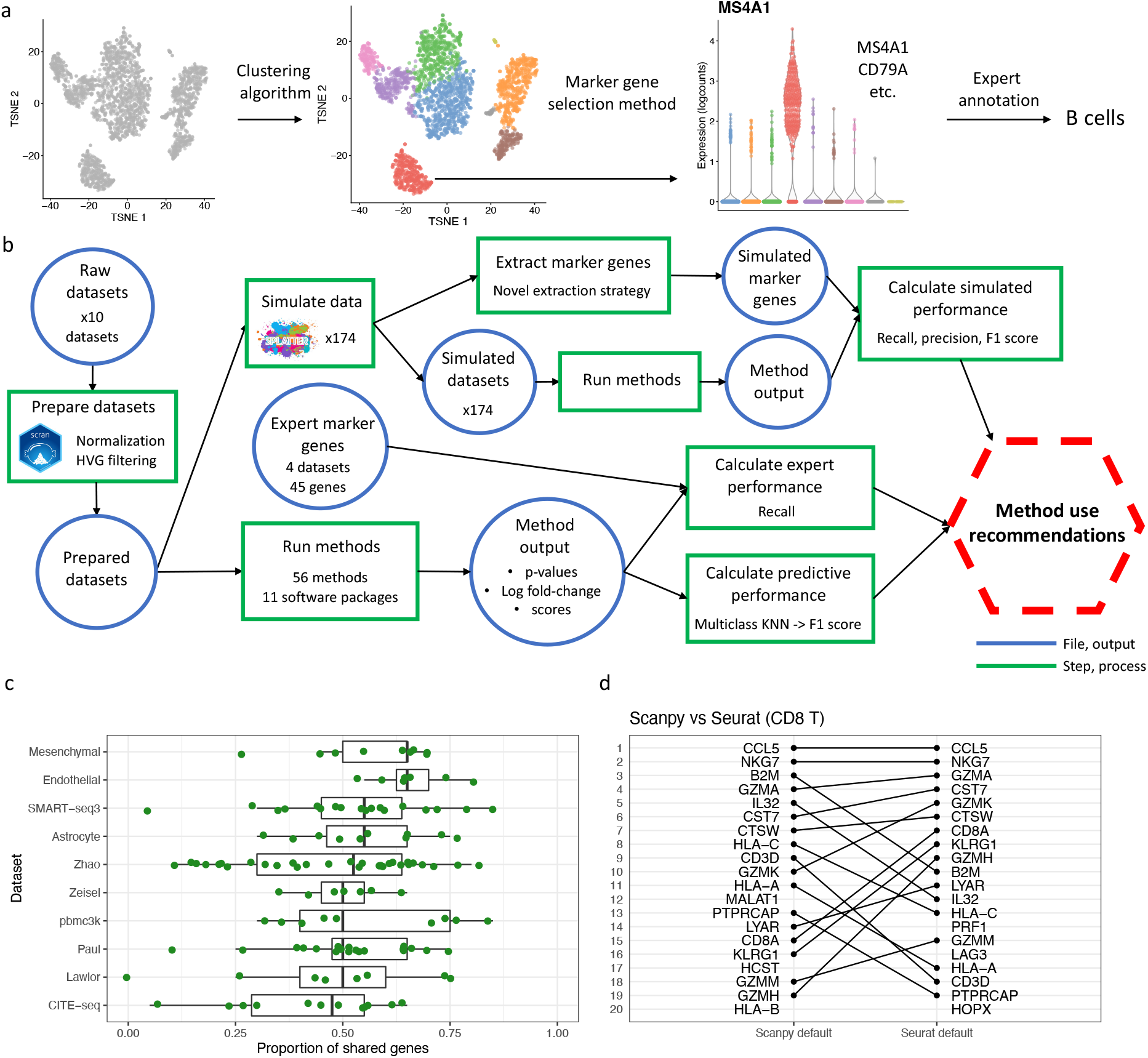
Overview of marker genes usage and benchmarking. a) A visual overview of the use of marker genes to annotate clusters. First, a clustering algorithm is performed to separate cells into putative clusters. Then, for each cluster, a marker gene selection method is used extract a small number of marker genes. This gene list is inspected and the expression of the genes visualised to give an expert-annotation of cell type for each cluster. b) A visual overview of the benchmarking performed in this paper. First, the real datasets are processed and the marker gene selection methods are run on the processed datasets. The output of the methods is extracted and used to calculate the methods’ predictive performance and ability to recover expert-annotated marker genes. The processed datasets are also used to simulate additional datasets, on which the methods are run and their ability to recover true simulated marker genes calculated. c) The proportion of shared genes in the top 20 genes selected by the default methods implemented by Scanpy and Seurat for each cluster across all 10 real datasets (127 clusters in total). d) A visual comparison of the rankings of the top 20 selected genes by the default Scanpy and Seurat methods in the CD8 T cell cluster in the pbmc3k dataset.

The concept of marker gene is closely related to that of a differentially expressed (DE) gene, but the two concepts are not synonymous. Broadly, we see marker gene, which refers to a gene that distinguishes a subpopulation, as a more abstract concept than a DE gene, which refers to a gene that shows a statistically significant difference in mean expression in a specific comparison. In the context of scRNA-seq data, the term DE gene does not have a unique meaning; it can refer to genes found in a comparison between cells in different clusters in the same sample ^[12,13]^ or in the same cluster between different samples ^[4,14,15]^, or to a comparison between arbitrary groups of cells^[16]^. When DE methods are used to identify marker genes a decision has to be made about how to map the idea of marker genes to a concrete between-groups comparison. Indeed, different methods use different concrete comparisons: Seurat and Scanpy use a ‘one-vs-rest’ cluster comparison strategy, while scran employs a ‘pairwise’ approach (see Methods for details). Significantly, both of these strategies are different to those compared in previous benchmarks of single-sample DE methods ^[12,17]^ that focused on comparing two specific cell subpopulations. For example, the ‘one-vs-rest’ strategy creates a situation with highly imbalanced sample sizes and increased biological heterogeneity in the pooled ‘other’ group, both of which we would expect to pose significant challenges to DE methods.

Today, many marker gene selection methods are available. These methods range from relatively simple differential expression based methods to those based on modern machine learning ideas. The most commonly used methods are implemented as part of the Seurat^[18]^ and Scanpy ^[19]^ analysis frameworks, which both implement a variety of method options. More recently, bespoke tools have also been developed, as marker gene selection continues to be an active area of method development ^[7,20,21]^. However, unlike other areas of scRNA-seq data analysis such as normalisation, differential expression analysis or trajectory inference ^[12,22,23]^, no impartial and comprehensive comparison of the different marker gene selection methods is available. The lack of such a comparison leaves analysts unaware of the relative performance of different methods. Furthermore, because the most commonly used methods are implemented in larger analysis frameworks, no publications exist that describe in detail or justify their marker gene selection methods. Therefore, their widely used methods are not supported by publicly scrutinised evidence of their effectiveness.

In this paper, we compared 56 methods for selecting marker genes in scRNA-seq data. The methods were benchmarked using 10 scRNA-seq datasets, encompassing a range of protocols and biological samples, as well as over 170 additional simulated datasets. Methods were compared on their ability to recover simulated and expert-annotated marker genes, the predictive performance and characteristics of the gene sets they select, their memory usage and speed, and their implementation quality (Figure 1b). To scrutinise the methodologies of the most commonly used methods, we created case studies based on the specific observations made during benchmarking. These case studies highlight several notable issues with, and inconsistencies between, the methods implemented in the Scanpy and Seurat packages.

Our comparisons and case studies demonstrated that while most methods perform well there is still variability in the quality of the marker genes selected. Significantly, more recent methods were not able to comprehensively outperform older methods. The case studies emphasised that there are large but unappreciated methodological differences even between similar methods, which can have large effects on their output in some scenarios. Overall, our results highlight the efficacy of simple methods, such as the Wilcoxon rank-sum test, for selecting marker genes.

## Results

### Scope of benchmarking

In this paper we use the phrase ‘marker gene’ as shorthand for ‘cell-subpopulation-specific marker gene’; that is, a gene whose expression distinguishes a particular cell subpopulation in a given dataset. As described above, marker genes can be used for a range of purposes, but in this paper we focus on their use in the annotation of the biological cell-type of clusters. This use of marker genes is by far their most common application in scRNA-seq data analysis. This focus, however, precludes an assessment of other types of methods that also select so-called marker genes. We do not consider methods that aim to select a small set of genes maximally informative for a given full clustering, rather than for each identified cluster ^[17,24,25]^. We also do not consider the ability of selected marker genes to be used in designing spatial transcriptomics analyses^[7,25]^. Finally, we only directly consider the quality of the marker genes in their use for annotation, rather than for other purposes, such as to gauge the quality of a clustering^[8]^ or to provide markers that can be used in wet-lab applications^[6]^. We believe, however, that methods that select high quality marker genes for annotation will likely also perform well at these other tasks.

### Method characteristics

The methods benchmarked in this paper differ in a number of important ways. First, they differ in the methodology they use to quantify the extent to which each gene can ‘distinguish’ cell types and is therefore a strong marker gene. Most methods use some form of differential expression (DE) testing (Seurat, Scanpy, scran `findMarkers()`, presto, edgeR, limma). Conversely, other methods use ideas of features selection (RankCorr), predictive performance (NSForest) or alternative statistics (Cepo, scran `scoreMarkers()`) to select marker genes. Within methods based on DE testing most (Scanpy, Seurat) use one-vs-rest testing while scran uses an alternative pairwise strategy (see Methods for details). In addition to differences in methodology there are substantive differences in the outputs of the methods: RankCorr and NSForest return only a specific set of genes determined to be marker genes, while other methods return a ranking of all genes for strength of marker-gene status, generally by p-value. While these methods do provide a strategy for selecting a set of marker genes, say taking those with a p-value below a given threshold, in practice the strategy is not feasible due to the small size of the returned p-values even after multiple testing correction (Figure S1, see below), which makes selecting a principled threshold challenging. To account for the difficulty of selecting a fixed set of markers we follow convention and select only a fixed-size set of the top *n* marker genes (for *n* = 5, 10, 20 say) as ranked by the method. In addition, another difference between methods is the direction of regulation of genes the method is designed to select. In this context a gene is up-regulated (down-regulated) if it is more (less) highly expressed in the cluster of interest relative to other clusters. Some methods, including Scanpy by default, will only select up-regulated marker genes, while other methods can select genes that are either up- or down-regulated (Figure 2b). Finally, a notable feature of the space of methods is duplication in methodology amongst the most commonly used Seurat, Scanpy and scran methods. We explore the relationship between Scanpy and Seurat methods in particular in case studies in this paper.

**Figure 2:**
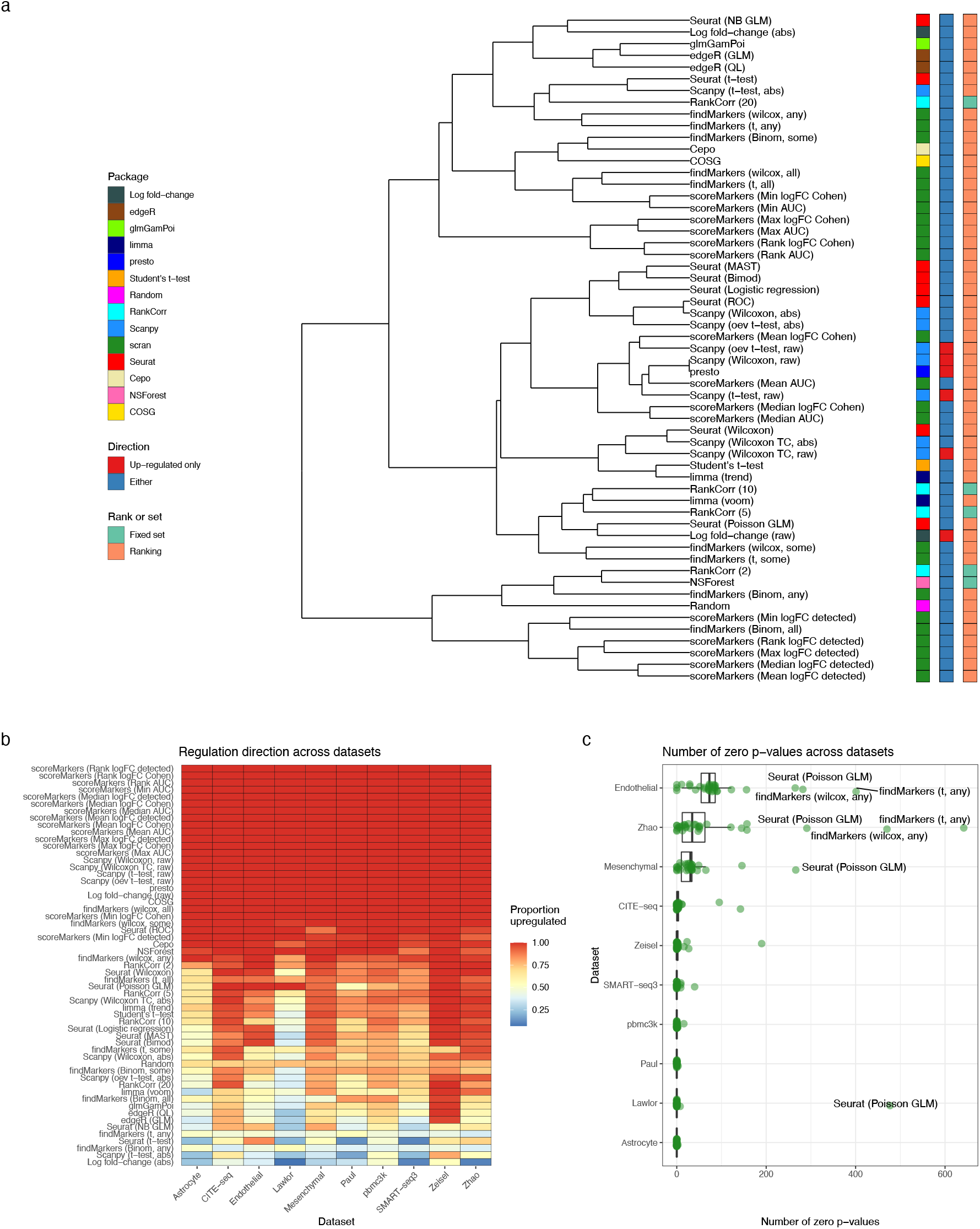
Method concordance and output characteristics. a) A dendrogram representation of the hierarchical clustering of methods based on the proportion of shared genes in the at most top 20 genes they select. Methods are labelled by the package which implements them, whether they select only up-regulated marker genes or both up- and down-regulated marker genes, and whether they output a set of marker genes or a ranking of gene by strength of marker gene status. b) The proportion of the at most top 20 genes selected of each method in each dataset (averaged over clusters). c) The number of genes for each method in each dataset (median over clusters) that returned p-values of exactly 0. Methods that did not return p-values were removed from this analysis.

### Benchmarking strategy

To compare the performance of the different methods we use a variety of metrics intended to provide a holistic assessment of a method’s performance. The diversity of techniques used by the methods to select marker genes means it is important that the metrics used to benchmark methods are not biased towards a particular set of techniques. For example, a simulation-based comparison could employ a similar generative probabilistic model to those used to perform statistical testing, potentially leading to overoptimistic conclusions. For this reason, we compare methods based on both their ability to recover simulated marker genes and the predictive performance of the gene sets they select. In addition, whether a gene is a marker gene depends on whether it is ‘known’ to be a marker gene^[26]^. Methods may be most useful if they find known marker genes, even if genes which better distinguish cell subpopulations are present in a dataset. To benchmark the methods’ ability to select ‘known’ marker genes we compare their ability to select expert-annotated marker genes in 4 datasets. In addition, we compare the methods based on their speed and memory efficiency and the quality of their software implementations Overall, we benchmarked the performance, computational efficiency, and output characteristics of 56 methods (Table 1) across 10 real datasets (Table 2) and over 170 additional simulated datasets. The real datasets were chosen to represent a range of protocols—including 10X Chromium, Smart-seq3, CITE-seq and MARS-seq—and a range of sizes, from approximately 3,000 to 60,0000 cells.

**Table 1:** Marker gene selection methods benchmarked in this paper

| Package | Version | Language | Parameters | Description | Citation |
| --- | --- | --- | --- | --- | --- |
| Seurat | 4.0.5 | R | test.use = t | Welch's t-test | [18] |
| - | - | - | test.use = "wilcox" | Wilcoxon rank-sum test (with tie-correction) | - |
| - | - | - | test.use = "LR" | Genewise Logistic regression | - |
| - | - | - | test.use = "negbinom" | Two group Negative Binomial GLM | - |
| - | - | - | test.use = "poisson" | Two group Poisson GLM | - |
| - | - | - | test.use = "roc" | ROC assessment of gene expression as classifier | - |
| - | - | - | test.use = "bimod" | Bimodal likelihood ratio test | -, [35] |
| - | - | - | test.use = "MAST" | MAST | -, [13] |
| COSG | 0.9.0 | R | None | Cosine score | [21] |
| Scanpy | 1.8.1 | Python | test.use = "t-test"<br>rankby.abs = True | Welch t-test<br>ranking by the absolute value of the score | [19] |
| - | - | - | test.use = "t-test.over.estimvar"<br>rankby.abs = True | Welch t-test with overestimated variance<br>ranking by the absolute value of the score | - |
| - | - | - | test.use = "t-test"<br>rankby.abs = False | Welch t-test<br>ranking by the raw score | - |
| - | - | - | test.use = "t-test.over.estimvar"<br>rankby.abs = False | Welch t-test with overestimated variance<br>ranking by the raw score | - |
| - | - | - | test.use = "wilcoxon"<br>rankby.abs = True<br>tie.correct = True | Wilcoxon rank-sum test<br>ranking by absolute value of the score<br>with tie-correction | - |
| - | - | - | test.use = "wilcoxon"<br>rankby.abs = False<br>tie.correct = True | Wilcoxon rank-sum test<br>ranking by the raw score<br>with tie-correction | - |
| - | - | - | test.use = "wilcoxon"<br>rankby.abs = True<br>tie.correct = False | Wilcoxon rank-sum test<br>ranking by absolute value of the score<br>without tie-correction | - |
| - | - | - | test.use = "wilcoxon"<br>rankby.abs = False<br>tie.correct = False | Wilcoxon rank-sum test<br>ranking by the raw score<br>without tie-correction | - |
| scran | 1.22.1 | R | findMarkers()<br>test.type = "t", pval.type = "any" | Pairwise t-test<br>up-ranking genes with "any" small p-values | [54] |
| - | - | - | findMarkers()<br>test.type = "t", pval.type = "all" | Pairwise t-test<br>up-ranking genes with "all" small p-values | - |
| - | - | - | findMarkers()<br>test.type = "t", pval.type = "some" | Pairwise t-test<br>up-ranking genes with "some" small p-values | - |
| - | - | - | findMarkers()<br>test.type = "wilcox", pval.type = "any" | Pairwise Wilcoxon rank-sum test<br>up-ranking genes with "any" small p-values | - |
| - | - | - | findMarkers()<br>test.type = "wilcox", pval.type = "all" | Pairwise Wilcoxon rank-sum test<br>up-ranking genes with "all" small p-values | - |
| - | - | - | findMarkers()<br>test.type = "wilcox", pval.type = "some" | Pairwise Wilcoxon rank-sum test<br>up-ranking genes with "some" small p-values | - |
| - | - | - | findMarkers()<br>test.type = "binom", pval.type = "any" | Pairwise Binomial test<br>up-ranking genes with "any" small p-values | - |
| - | - | - | findMarkers()<br>test.type = "binom", pval.type = "all" | Pairwise Binomial test<br>up-ranking genes with "all" small p-values | - |
| - | - | - | findMarkers()<br>test.type = "binom", pval.type = "some" | Pairwise Binomial test<br>up-ranking genes with "some" small p-values | - |
| scran | 1.22.1 | R | scoreMarkers(), mean.logFC.cohen | Pairwise Cohen's d, mean across genes | NA |
| - | - | - | scoreMarkers(), min.logFC.cohen | Pairwise Cohen's d, minimum across genes | - |
| - | - | - | scoreMarkers(), median.logFC.cohen | Pairwise Cohen's d, median across genes | - |
| - | - | - | scoreMarkers(), max.logFC.cohen | Pairwise Cohen's d, maximum across genes | - |
| - | - | - | scoreMarkers(), rank.logFC.cohen | Pairwise Cohen's d, min rank across genes | - |
| - | - | - | scoreMarkers(), mean.AUC | Pairwise AUC, mean across genes | - |
| - | - | - | scoreMarkers(), median.AUC | Pairwise AUC, median across genes | - |
| - | - | - | scoreMarkers(), min.AUC | Pairwise AUC, minimum across genes | - |
| - | - | - | scoreMarkers(), max.AUC | Pairwise AUC, maximum across genes | - |
| - | - | - | scoreMarkers(), rank.AUC | Pairwise AUC, min rank across genes | - |
| - | - | - | scoreMarkers(), mean.logFC.detected | Pairwise lfc in detection proportion, mean across genes | - |
| - | - | - | scoreMarkers(), min.logFC.detected | Pairwise lfc in detection proportion, minimum across genes | - |
| - | - | - | scoreMarkers(), median.logFC.detected | Pairwise lfc in detection proportion, median across genes | - |
| - | - | - | scoreMarkers(), max.logFC.detected | Pairwise lfc in detection proportion, maximum across genes | - |
| - | - | - | scoreMarkers(), rank.logFC.detected | Pairwise lfc in detection proportion, min rank across genes | - |
| presto | 1.0.0 | R | None | Optimised Wilcoxon rank-sum test | NA |
| edger | 3.36.0 | R | GLM | Negative Binomial GLM with empirical Bayes (EB) shrinkage | [62,63] |
| edger | 3.36.0 | R | QL | - with quasi-likelihood fitting | [62,63] |
| RankCorr | NA | Python | lambda = 2 | Sparse separating hyperplane selection | [26] |
| RankCorr | NA | Python | lambda = 5 | - | [26] |
| RankCorr | NA | Python | lambda = 10 | - | [26] |
| glmGamPoi | 1.6.0 | R | None | Negative Binomial QL GLM with EB shrinkage | [64] |
| limma | 3.50.0 | R | standard | Linear model with EB shrinkage | [65,66] |
| limma | 3.50.0 | R | voom | - with weighting | [65,66] |
| limma | 3.50.0 | R | trend | - | [65,66] |
| Cepo | 1.0.0 | R | None | Stability statistic | [67] |
| NSForest | 3.0 | Python | None | Random forest classifier | [20] |

**Table 2:** Real scRNA-seq datasets used in this paper

| Dataset | N. cells | N. cell populations | Description | Technology | Citation |
| --- | --- | --- | --- | --- | --- |
| Zeisel | 3005 | 7 | Mouse brain | 10X Chromium | [55] |
| pbmc3k | 2638 | 9 | PBMC | 10X Chromium | [61] |
| Lawlor | 604 | 8 | Human pancreas | SMARTer v1 | [32] |
| Endothelial | 30641 | 7 | Human lung endothelium | 10X Chromium | - |
| Astrocyte | 3013 | 10 | Mouse brain astrocytes | Single-nucleus RNA-seq | [8] |
| Paul15 | 2730 | 19 | Mouse hematopoietic stem cells | MARS-seq | [68] |
| Zhao | 60672 | 29 | Human liver resident immune cells | 10X Chromium | [69] |
| SS3 PBMC | 3128 | 17 | PBMC | Smart-seq3 | [56] |
| CITE-seq | 8617 | 12 | CBMC | CITE-seq | [70] |
| Mesenchymal | 11408 | 9 | Human lung mesenchymal | 10X Chromium | - |

### Method concordance and selected marker gene characteristics

First, we compared the extent to which the different methods produced concordant gene sets. To motivate this analysis, and the benchmarking more broadly, we compared the concordance of the default methods implemented by Scanpy and Seurat, which are the two most commonly used methods. For the top 20 genes selected by the methods we calculated the proportion of genes shared between the methods and visualised the difference in their rankings. Despite their similar methodology the methods showed only poor to moderate concordance. Across datasets many clusters showed less than 50% overlap between the two methods (Figure 1c). Even in clusters with a high proportion of shared genes, such as the CD8 T cluster in the pbmc3k dataset, the methods’ rankings were quite different (Figure 1d).

To examine the concordance of all the methods more generally we performed a hierarchical clustering based on the proportion of genes shared between methods averaged across clusters and datasets (Figure 2a; see Methods for details). This analysis recapitulated many known features of the space of methods: the dendrogram groups together similar methodologies implemented in different packages and methods that only select up-regulated genes (Figure 2). In addition, the clustering also revealed that some Seurat methods with different methodologies give similar rankings.

Next, we examined the characteristics of the marker genes produced by the analysis. First, we calculated the proportion of the (at most) 20 top genes for each method that are up-regulated, averaging over clusters (Figure 2b). In this context, up-regulated means that the gene is more highly expressed in the cluster of interest than in other clusters. Generally, marker genes are up-regulated. For example, all of the 45 expert-annotated marker genes we consider in this paper are up-regulated. Most methods select a majority of up-regulated genes. A subset of methods including presto, scran’s `findMarkers()` Wilcoxon and `scoreMarkers()` methods, and the Scanpy methods only selected up-regulated genes, while Welch t-test and Binomial test based methods select mainly down-regulated genes. In addition, for the DE-based methods we extracted and visualised p-values calculated by the methods. As noted above, the p-values were extremely small even after conventional multiple testing correction (Figure S1). The small magnitude of the p-values is caused by the large sample sizes in the datasets (where each cell is treated as a replicate) and double dipping inherent in performing clustering and statistical testing on the same data (Figure S2). Surprisingly, many methods showed p-values of exactly 0, especially in the larger Zhao, Endothelial and Mesenchymal datasets (Figure 2c). This feature occurred for methods implemented in both R and Python, and has a large impact on Seurat’s methods as described in the case studies section.

### Performance on simulated marker genes

To critically assess the performance of the methods we first compared their ability to recover true simulated marker genes. We use the splat simulation model implemented in the splatter R package to simulate scRNA-seq datasets^[27]^. The splat model is the most widely used simulation model in scRNA-seq benchmarking studies^[28]^ and was recently shown to be in the top tier of simulation methods^[29]^. To simulate marker genes specifically we designed and implemented a novel marker gene score to select and rank marker genes using the splat model’s cluster-specific differential expression parameters (Methods). A limitation of the splat model is that it does not allow these cluster-specific differential expression parameters to be estimated from data. Indeed, no current simulation scRNA-seq method incorporates this facility^[28]^. To overcome this limitation in practice we performed analyses comparing simulated and true marker genes to identify values of the parameters that were able to recapitulate true expert-annotated marker genes (Figure S8, Figure S7: compare with e.g., Figure S4, Figure S5). Our simulations also include sensitivity analyses using a range of plausible values for these parameters.

Ten simulation scenarios were considered, each with parameters estimated from one of the real datasets (Table 2). All simulations used to compare performance had 2,000 genes, 2,000 cells, and 5 clusters. In this analysis we only simulate up-regulated marker genes and the outputs of the methods are also filtered to include only up-regulated marker genes (see Methods for details). Each simulation scenario was repeated 3 times to gauge variability. For each method in each simulated cluster we select the top 20 simulated marker genes and the (at most) top 20 marker genes selected by each method. With the top sets of genes we computed the precision, recall, and F1 score of each method in each cluster and replicate (Figure S11). This analysis showed little replicate- and cluster-specific variability so we summarised the performance of each method in each scenario by taking the median across clusters and replicates (Figure 3).

**Figure 3:**
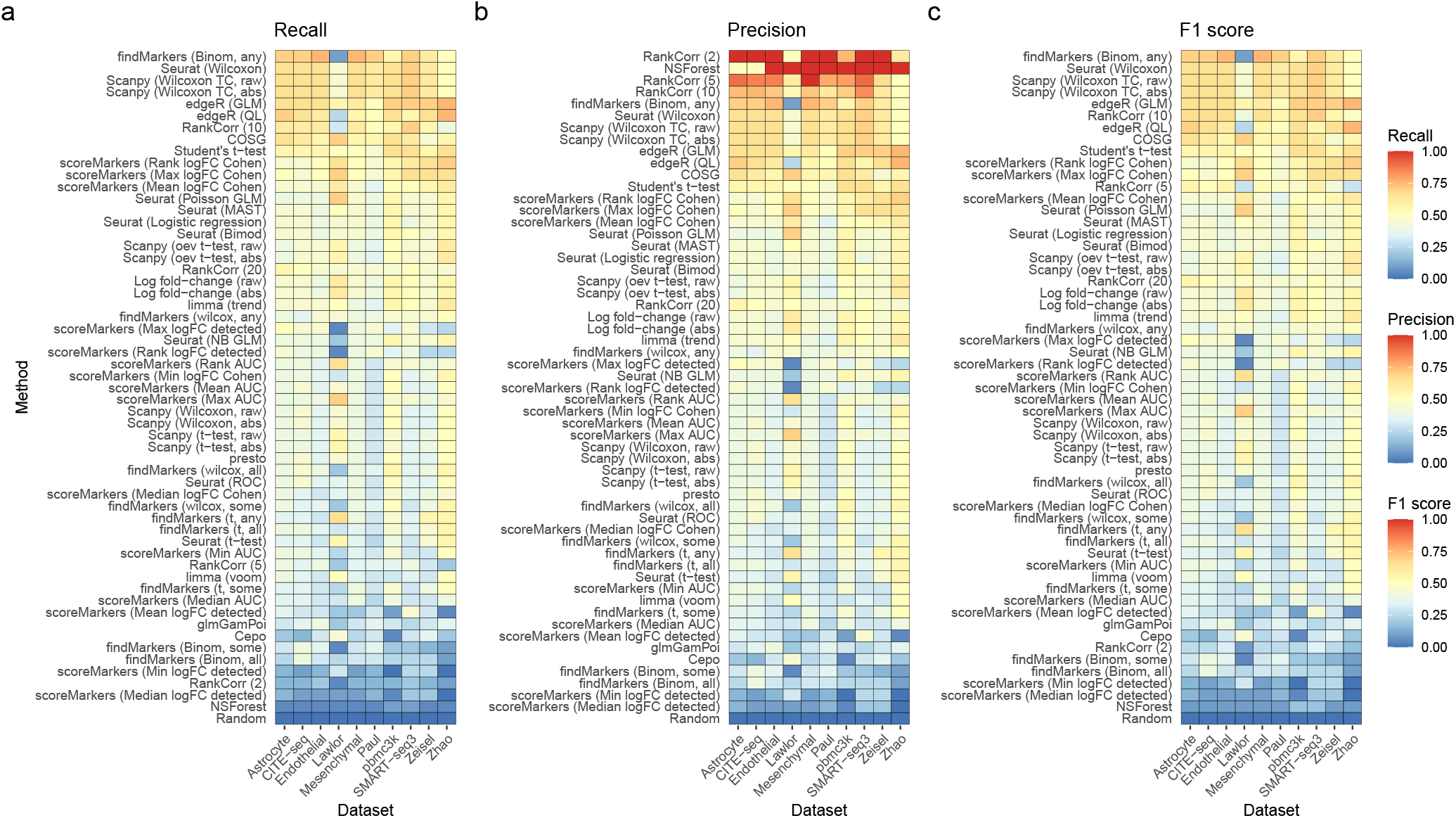
Comparison of methods using simulated datasets. a) Calculated recall for all methods on simulations scenarios based on all real datasets. The marker genes selected based on the simulation model parameters are used as the ground truth. Methods are ranked top to bottom in the heatmap by median recall across scenarios. b) As in a) but now with precision. c) As in b) but calculating F1 score. These results average over simulated clusters and simulation replicates and are conducted with 20 genes selected and a location parameter for the DE factor in the splat simulation model of 3.

Overall, we found concordant results across simulation scenarios. Assessed by recall (the proportion of all true marker genes that are selected as marker genes by the method), the best performing methods were the scran’s Binomial-any method, Wilcoxon rank-sum based methods (Seurat and Scanpy), and edgeR methods, as well as Student’s t-test and RankCorr with the lambda parameter set to 10. The worst methods were NSForest, Cepo and RankCorr (2) (Figure 3a). Despite the strong performance of scran’s Binomial- any method the rest of scran’s methods showed relatively weak performance. Assessed by precision (the proportion of marker genes a method selects that are true marker genes), the best performing methods were RankCorr and NSForest as well as scran Binomial-any (Figure 3b). The discrepancy between precision and recall for RankCorr and NSForest occurs because RankCorr and NSForest only select a very small number of genes. To provide an overall ranking of the methods we calculated the F1 score (a combination of recall and precision) for each method and dataset and then ranked the methods by their median performance over datasets (Figure 3c). In this ranking, the best performing methods were the scran Binomial-any method, the Wilcoxon rank-sum based methods and edgeR’s methods. NSForest, Cepo and scran’s other Binomial methods showed the worst performance. While the results were generally concordant between scenarios the Lawlor dataset-based scenarios showed some differences: here the log fold-change methods and Seurat’s Poisson GLM were more effective, while scran Binomial-any was less effective. This difference is likely caused by the higher mean counts of the Lawlor dataset relative to other datasets (Figure S3).

Further analyses assessed the sensitivity of these results to different analysis parameters. Using different values (*n* = 5, 10, 20, 40) for the number of genes had little effect on the overall ranking, though scran Binomial-any showed worse, and edgeR improved, performance when 40 genes were used (Figure S10). Using different values of the cluster-specific differential expression parameters also had little effect, although the COSG method did perform better when smaller values were used.

### Expert-annotated marker gene recovery

Next, we compared the ability of the methods to recover expert-annotated sets of marker genes. This analysis is intended to assess the ability of methods to select genes which are *known* marker genes. We used expert-annotated marker gene sets for four datasets: the Lawlor, Smart-seq3, pbmc3k and Zeisel datasets, derived mainly from the papers and tutorials describing these datasets (see Methods for details). Initially, we hoped to perform a comparison based on genes available in online databases such as CellMarker^[30]^ and PangloDB^[31]^, however we found that there were substantial differences between databases, and that marker-gene coverage for specific cell types was typically sparse. In addition, it was difficult to harmonise cell-type labels between the databases and our specific datasets, especially for the PBMC datasets. For these reasons we instead used sets of marker genes created with respect to the specific datasets.

Concretely, for each method and cluster we computed recall using the expert-annotated marker genes as the ground truth. Furthermore, we calculated the number of clusters that could be successfully annotated using the method’s selected marker genes. For this calculation we considered a cluster to be successfully annotated if all its associated expert-annotated marker genes were selected by the method. In these analyses, the number of selected marker genes used was based on the number of expert-annotated marker genes per cluster. First, for the Lawlor dataset, we used the canonical markers used for classification in the paper describing the dataset^[32]^ and selected the top gene for each method. In this dataset, Seurat’s ROC, logistic regression, Poisson GLM and Bimod methods, presto, limma, ranking by the raw log fold-change, Scanpy’s Wilcoxon methods, `scoreMarkers()`’s min-cohen, min-AUC and mean-AUC methods. and Student’s t-test successfully selected all expert-annotated marker genes while again scran’s `findMarkers()` methods and Cepo showed poor performance (Figure 4a, c). Second, for the pbmc3k dataset we used the marker genes given in the Seurat Guided Clustering Tutorial^[33]^ and used the (at most) top 10 selected genes. In this dataset, the best performing methods were the log fold-change (raw), and Student’s t-test, while the worst performing methods were the t-test-based scran methods, Cepo and RankCorr (Figure 4b, d). For example, the Student’s t-test selected marker genes could be used to annotate 7 clusters, while those selected by Cepo could only annotate 2 (Figure 4d). Strikingly, all methods struggled to select the expert-annotated marker genes for the Memory and Naive CD4 T cell clusters. This effect occurrs because the expert-annotated marker genes for these clusters, especially *IL7R*, have expression profiles that separate all T cells from other clusters, not the specific T cell subtypes of interest (Figure S6).

**Figure 4:**
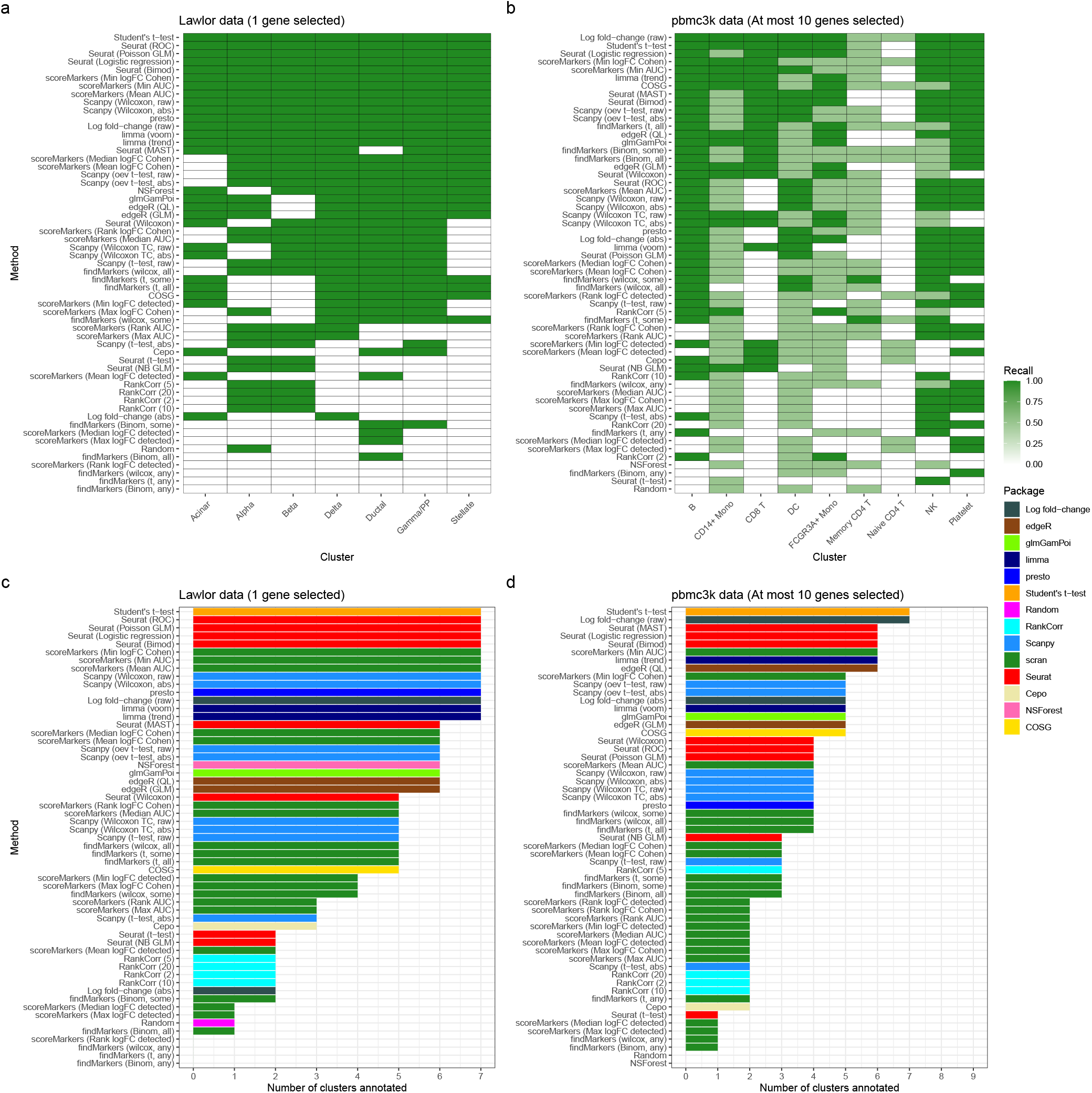
Comparison of methods based on expert-annotated marker genes. a) Recall of methods when selecting marker genes in the Lawlor dataset using a set of expert-annotated marker genes as the ground truth. The marker genes used to annotate the clustering in the original publication describing the Lawlor dataset were used as the set of expert-annotated marker genes. The top gene was selected from the output of each method. b) As in a) but for the pbmc3k dataset, using the (at most) top 10 marker genes from each method and taking the set of expert-annotated marker genes to be those used in the Seurat package’s “Guided Clustering Tutorial”. c) The number of clusters that are successfully annotated using the selected marker genes in the Lawlor datasets (other details as in a)). A specific cluster is defined as successfully annotated if the selected marker genes include all the expert-annotated marker genes for that cluster. d) The same success of annotation analysis as in c) but for the pbmc3k dataset, with the details of the expert-annotated marker genes and number of selected marker genes as in b).

Similarly, we calculated the performance of the methods on the expert-marker genes for the Smart-seq3 and Zeisel data. In these comparisons the relative performance of the methods was similar to the pbmc3k and Lawlor datasets but the absolute performance of methods was generally worse, although some scran methods did show improved performance in the Smart-seq3 dataset (Figure S12, Figure S13). Examination of the results suggested that the poor absolute performance was mainly due to marker genes being difficult to select or because the methods selected other strong marker genes. For example, *PECAM1*, the marker of Naive/Memory CD8 T cells in the Smart-seq3 PBMC dataset, is not found because its expression profile means that it is only a good marker relative to other T cells and not globally, which is the implicit frame of reference of the methods (Figure S14). On the other hand, in the Zeisel dataset’s oligodendrocyte cluster Seurat’s Wilcoxon method’s top 10 selected marker genes are all strong markers, but the expert-annotated marker gene *HAPLN2* is simply not selected (Figure S4). In addition, examining the genes selected by poor performing methods for these datasets reveals reasons for their poor performance. For the interneuron cluster of the Zeisel dataset the top 2 genes selected by Seurat’s t-test method are strongly up-regulated in the interneuron cluster but are also up-regulated in most other clusters (Figure S5a). In contrast, the top 2 genes selected by scran’s t-test-any method for the interneuron cluster of the Zeisel dataset show strong up-regulation only in a different (i.e. not interneuron) cluster, making them poor marker genes of the interneuron cluster (Figure S5b).

### Predictive performance

To gain a different perspective on marker gene selection, we compared the predictive accuracy of the gene sets selected by competing methods. The idea of this comparison is that better gene sets should capture more ‘information’ about which cells are in each cluster. We quantify the amount of information by measuring the predictive performance of a classifier for (multi-class) cluster status trained on only the gene sets selected by the methods. Specifically, we select the top 5 marker genes for each method, dataset, cluster combination. The genes for different clusters are then pooled to give an overall gene set for each dataset. Using only the data for these genes a KNN classifier (with three nearest neighbours) or an SVM classifier (with linear kernel) are trained to predict the assigned cell-type of the cells. We found that the results from the KNN classifier (Figure 5) and the SVM classifer (Figure S15) were extremely similar (except for the unusual Lawlor dataset Figure S3) so we present only the results for the KNN classifier in the main text. To assess the variability in performance k-fold cross validation with 5 folds was used. While conceptually entirely distinct from other comparison methods applied above, this comparison was able to reproduce features seen in other comparisons, such as the difficulty of distinguishing between the Naive CD4 T and Memory CD4 T cells in the pbmc3k dataset (Figure 5a). To assess the performance of the methods we calculated a median F1 score across clusters for each fold and dataset (Figure 5b). Overall, the fold-based variability in each method’s performance was relatively low. To summarise each method’s performance across datasets we computed the z-score transformation of F1 score within datasets and then ranked the methods by their mean z-scores (lowest first) across datasets (Figure 5c). Using z-scores accounts for the wide variability in F1 scores observed across datasets, likely due to differences in the quality of their clusterings. Across datasets the best performing methods are the Seurat logistic regression, MAST and bimod methods, Student’s t-test, raw log fold-change ranking and RankCorr (5), while the worst performing methods included Cepo, Seurat’s t-test method, NSForest, absolute value log fold-change ranking, and scran’s binomial test methods. The Wilcoxon rank-sum test methods in Scanpy and Seurat also perform well.

**Figure 5:**
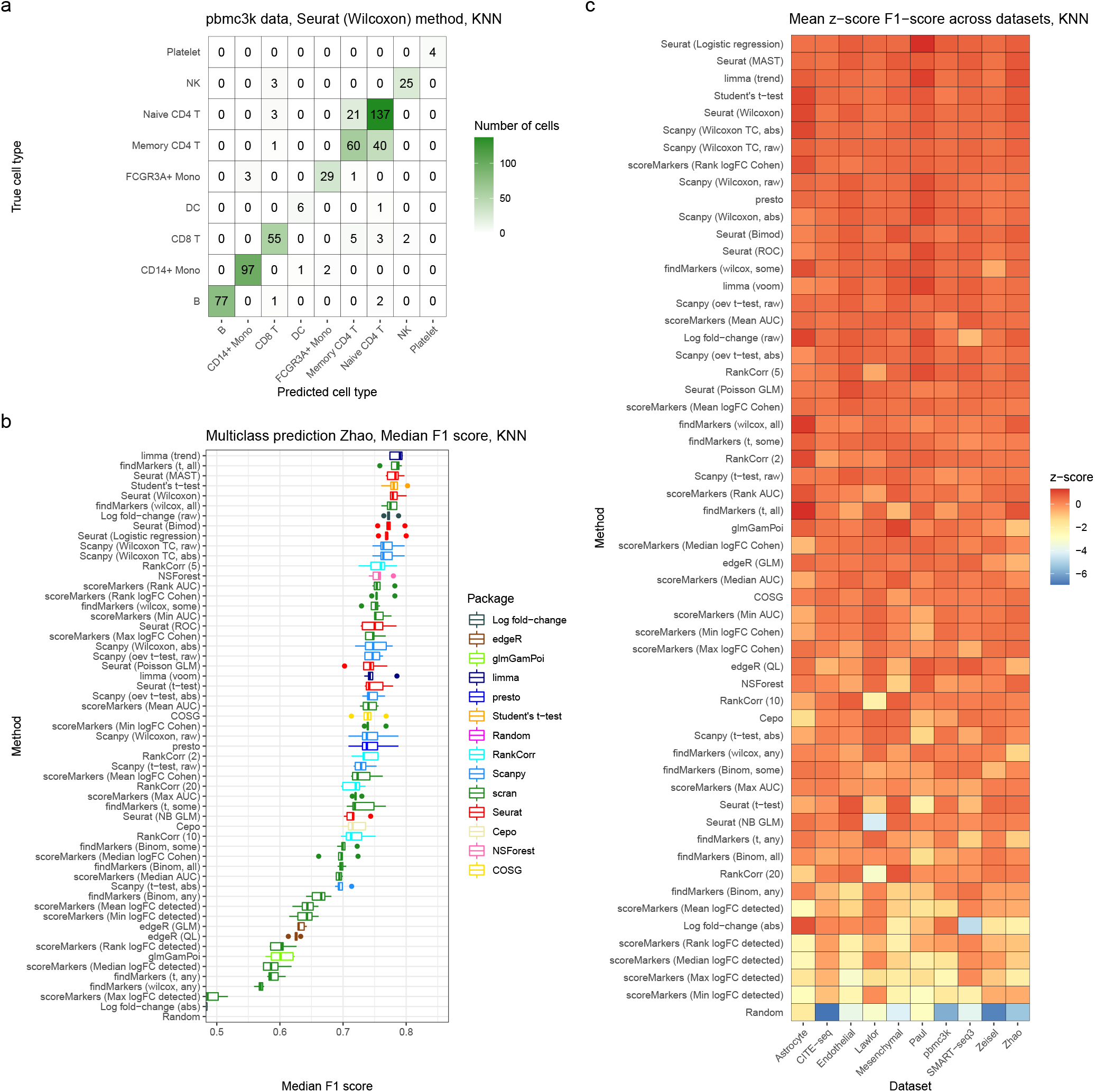
Comparison of methods using predictive performance. a) A confusion matrix representation of the performance of a KNN (three nearest neighbours) classifier using the set of genes selected by the Seurat Wilcoxon method in the pbmc3k dataset. b) Median F1 score of the KNN classifier using genes selected by all marker gene selection methods in the Zhao dataset. Each point is the F1 score in one of the 5 folds. c) The z-score of the median F1 score of the KNN classifier (averaging across folds) in each dataset. Methods are ranked top to bottom by their mean z-score across datasets.

### Computational performance and implementation quality

Finally, we compared the computational performance of the methods and the quality of their implementations. First, the methods were compared based on their speed and memory consumption (see Methods for details of how these were measured). Speed is particularly important for marker gene selection methods given the common need to run them multiple times when iterating through different clusterings of the data. The methods were run on all datasets as well as additional simulated datasets where the total number of cells and number of clusters were varied. Running the methods on all datasets highlighted that the methods had a very wide range of speeds (Figure 6a). For example, on the large Zhao dataset (60,672 cells) edgeR runs in over 10 hours while Scanpy’s default method runs in less than 15 seconds. Overall, the slowest methods were the edgeR methods, Seurat’s NB GLM and MAST methods, and NSForest, while the fastest were most of Scanpy’s methods, presto, Cepo and RankCorr. Furthermore, we observed that Seurat’s methods are substantially slower than scran and Scanpy’s methods even when implementing the same statistical tests. For example, on the Zhao dataset Seurat’s t-test method took over 90 minutes to run, while the equivalent Scanpy method took 14 seconds. These results were strongly concordant across simulations with varying numbers of total cells and number of clusters, which showed little change in the ordering of methods as the number of cells or clusters increased (Figure 6c, Figure S17a). Next, we compared the memory usage of all methods across datasets (Figure 6b). These measurements showed that memory usage was substantially higher across methods for the larger datasets (Endothelial, Mesenchymal, Zhao) studied. The edgeR, limma (voom) and glmGamPoi methods used the most memory, while scran methods and presto used the least. Simulations with varying numbers of total cells and clusters particularly highlighted the high memory usage of edgeR, glmGamPoi and limma when the total number of cells was high (Figure S16, Figure S17b)

**Figure 6:**
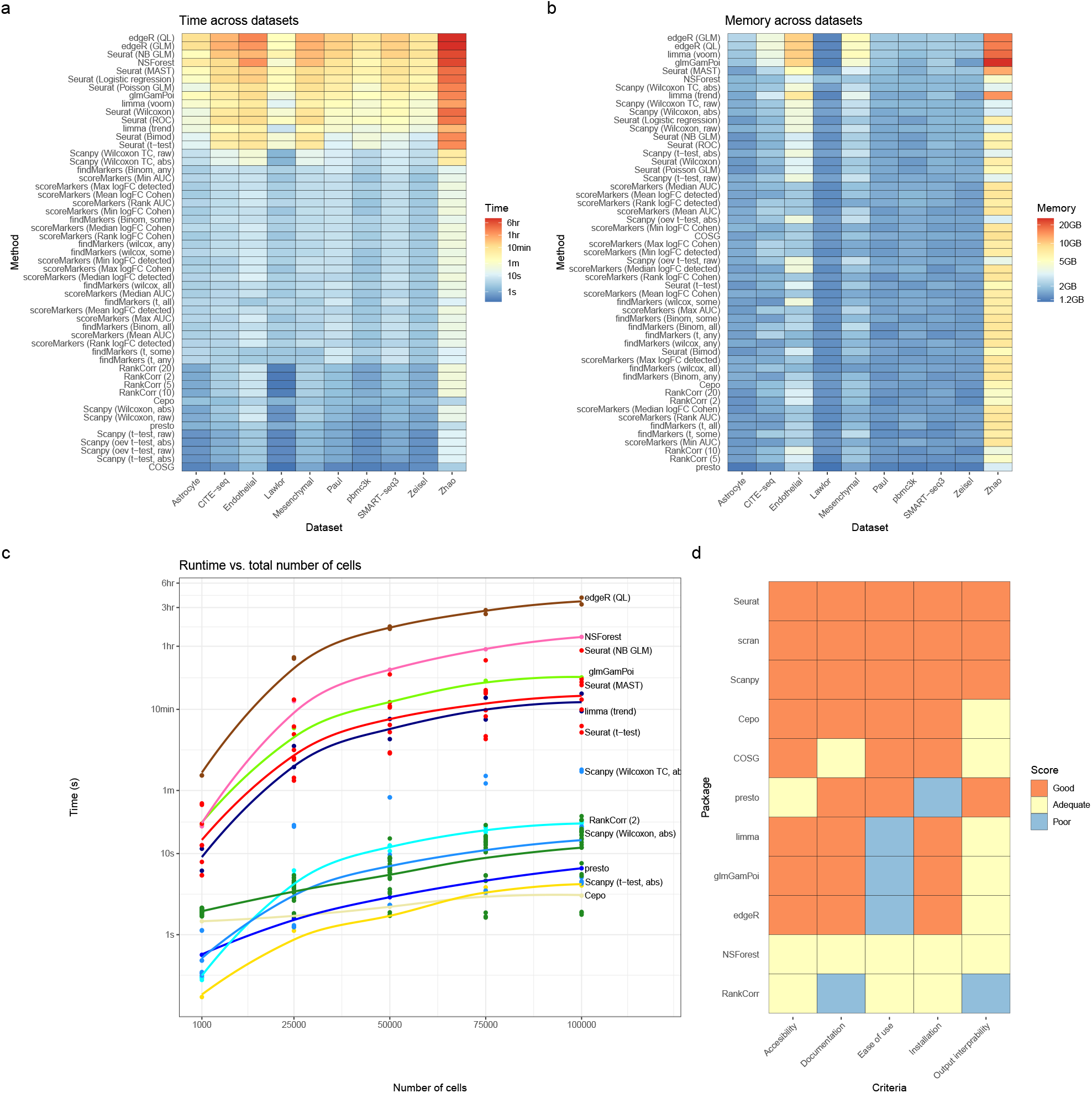
Comparisons of methods computational performance and implementation quality. a) Heatmap displaying the time taken for all methods to run across all 10 real datasets. Methods are ranked top to bottom in the heatmap by median time taken over dataset, largest at the top. Note that the colour scale of the heatmap is a log scale. b) Heatmap displaying the memory usage of all methods across all datasets. Methods are ranked top to bottom in the heatmap by median memory usage over datasets, largest at the top. c) Time taken for all methods on simulated datasets with increasing numbers of total cells. Points are averages over 3 simulation replciates. All simulations had parameters estimated from the pbmc3k dataset and a location parameter for the DE factor of 3. d) Assessment of the implementation quality of packages which implement methods for selecting marker genes based on 5 criteria.

For a final assessment of usability we compared the quality of the software implementing each method. Although this analysis is somewhat subjective, how easy a method is to install and run is one of the strongest determinants of its practical utility. For this analysis we consider methods on the level of packages. To compare the packages we assessed their performance as either good, adequate or poor for five criteria: accessibility, installation, documentation, ease of use and the interpretability of output (Figure 6d; see Methods for details). The Seurat, Scanpy and scran packages have excellent implementations, while Cepo is overall very good expect for difficult-to-use output that mangles cluster names. The COSG method has a good implementation, though with somewhat limited documentation and output interpretability. Next, edgeR, limma, glmGamPoi and presto all have good implementations with particular issues: the general purpose edgeR, limma and glmGamPoi methods need additional code to be used to select marker genes, while presto is difficult to install and is not available from standard repositories. Finally, NSForest and RankCorr are implemented only in the form of Python scripts in GitHub repositories with little documentation.

### Case studies

During the overall benchmarking we noticed several undocumented features of the methods implemented in Scanpy and Seurat, which can lead both to inconsistencies between Scanpy and Seurat and substantive issues with their methods’ output.

The major differences between Seurat and Scanpy’s methods are the strategies they use to rank genes after differential expression testing has been performed. Seurat ranks genes based first on their corresponding p-value (smallest first) and secondly (i.e., to break ties in the p-values) by the raw value of the log fold-change calculated for each gene (largest first). On the other hand, Scanpy ranks genes based on a test statistic or ‘score’ used in the calculation of the differential expression for each gene. By default, genes are ranked by the raw value of the statistic, such as a z-score, but they can be optionally ranked by the absolute value of the statistic. For the statistics implemented in Scanpy the sign of the statistic is the same as the log fold-change calculated for each gene.

These differences cause several issues. First, it creates a situation whereby the two methods select qualitatively different sets of marker genes: as noted above Scanpy by default only selects marker genes that are up-regulated, while Seurat by default can select both up- and down-regulated marker genes (Figure 2b). While both options are sensible, it is unlikely that many analysts recognise this difference or check that default method behaviour matches their expectations for marker gene selection.

Next, returning many p-values exactly equal to zero, as observed to occur for large datasets (Figure 2c), leads to an unexpected and undocumented change in the output of Seurat’s methods. The ranking of genes with zero p-values, of which there can often be more than the number of marker genes selected, is completely determined by the gene’s log fold-change value. (Figure 7a). Therefore, for large datasets, the output from Seurat’s methods will be the same as ranking by the (raw) log-fold change, while Scanpy’s methods are different (Figure 7b).

**Figure 7:**
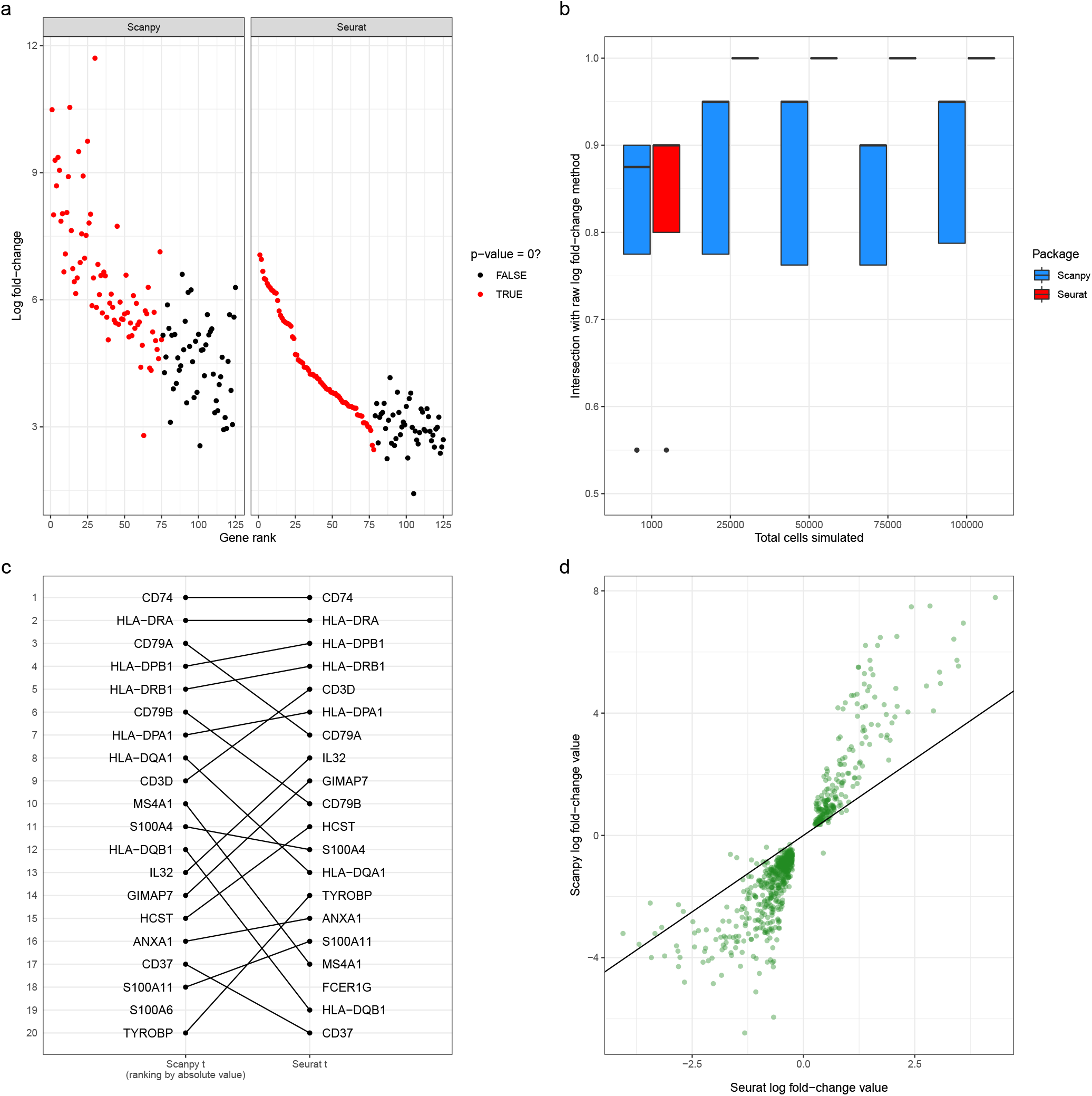
Case studies scrutinising Scanpy and Seurat. a) Gene rank vs log fold-change values for the Scanpy Wilcoxon (with tie correction, ranking by the absolute value of the score) and Seurat Wilcoxon methods for the Oligodendrocyte cell type cluster in the Zeisel dataset. The colour of the point indicates whether or not the genes has an exactly zero p-value. b) The proportion of top 20 genes shared between methods implemented in Scanpy and Seurat and the method of ranking genes by the raw log fold-change calculation on simulated datasets with increasing number of total cells. These simulations have parameters estimated from the pbmc3k dataset and a location parameter for the DE factor of 3. c) Visualisation of difference in rankings between the Scanpy t method (ranking by the absolute value of the score) and Seurat’s t-test method on the B cell cluster in the pbmc3k dataset. d) Scatter plot between the log fold-change values calculated by Seurat and Scanpy on the B cell cluster in the pbmc3k dataset.

Finally, Scanpy’s ranking by score means that its default methods output a mathematically incorrect ranking of genes. Scanpy uses both a z-score (Wilcoxon rank-sum test) and a t-value (Welch’s t-test) to rank genes. Due to the monotonicity of the cumulative density function, ranking by the absolute value of the z-score is equivalent to ranking by the p-value (see Methods). However, this equivalence does not hold for the t-value calculated from Welch’s t-test (Methods). This difference can be seen when comparing the output between Seurat and Scanpy Welch t-test methods: some genes, including those in the top 5, change their rankings substantially (Figure 7c).

In addition, there are other differences between Scanpy and Seurat, beyond their ranking strategies. First, by default they use different versions of the Wilcoxon rank-sum test. By default, Scanpy does not implement the conventional correction for ties (although it will with certain options set), while Seurat does. Second, a more serious difference is that the methods calculate different values for the same gene’s log fold-change in the same dataset (Figure 7d). This discrepancy is caused by the methods using different formulas to calculate the log-fold change values (see Methods for details). Extreme care should be taken when comparing the log fold-changes output by Seurat and Scanpy.

Finally, both methods produce spurious results when a gene has exactly zero expression (i.e. zero counts in all cells) in a cluster. When a gene has zero expression, Scanpy’s calculated log-fold change values become spuriously large (Figure S18a), while Seurat’s t-test method will preferentially up-rank genes that have zero expression even if better candidate marker genes are available (Figure S18b).

## Discussion

Marker gene selection is a ubiquitous component of the analysis of scRNA-seq data. However, despite its importance and the large number of methods available, to date there has been no impartial and systematic comparison of competing methods. In this paper, we provided the first comprehensive benchmarking of the available methods. We compared the methods’ performance on a range of metrics and scrutinised the most commonly used methods implemented in the Scanpy and Seurat analysis frameworks. While related to previous work comparing methods for selecting between-cluster DE genes^[12]^, our work focused explicitly on the challenges of finding marker genes and used both a different simulation setting and other different metrics to compare methods.

Overall, across comparisons the best performing methods were those based on the Wilcoxon rank-sum test, Student’s t-test or logistic regression. Our results also highlighted that several methods showed systematically worse performance: scran’s `findMarkers()` methods (especially using the default p-value any method), Cepo, and Welch t-test based methods when both up- and down-regulated genes are selected. In addition, while methods that only selected a subset of genes (RankCorr and NSForest) had excellent specificity they did not show strong predictive performance. Our results also highlighted large differences between the speed and memory consumption of different methods: methods designed for bulk RNA-seq data were particularly memory intensive, while Seurat’s methods were unexpectedly slow. Taken together, these results highlight that, in general, more complex methods are not required to select marker genes in scRNA-seq data.

Furthermore, our scrutinisation of Scanpy and Seurat highlighted issues and inconsistencies with their implemented methods. Most significantly, we found that due to an interaction between the surprising existence of zero p-values and Seurat’s ranking strategy most of Seurat’s methods will give identical results to ranking by the raw log fold-change values when the number of cells is large. In addition, we identified inconsistencies between how Seurat and Scanpy calculate the log fold-change values that make comparing their output fraught. The existence of these issues highlights the challenges of implementing large analysis frameworks which implement many methodological details. While such frameworks are extremely useful for analysts and contribute greatly to the field, the size of their impact magnifies the downstream effects of any issues in their implementations.

The analysis in this paper suggests a number of directions for further research into improving methodology for selecting marker genes. First, it suggests that attention should be refocused on improving the performance of simple Wilcoxon rank-sum and Student’s t tests, which show strong performance even without substantial adaption to the problem of marker genes. Second, our analysis of the DE-based methods’ p-values highlight the need for more extensive work on correcting p-values for double-dipping. Such work would enable marker genes to be selected based on well-calibrated p-value thresholds rather than an ad-hoc number of genes cutoff. Recent work on post-selection inference after clustering represents a promising direction for addressing this issue^[34]^. Third, specific examples in our analysis highlight that no methods can select marker genes at multiple levels of resolution, for example finding marker genes for both all T cells and specific T cell subsets. Selecting multi-resolution marker genes, likely in tandem with advances in multi-resolution clustering, would allow more precise characterisation of cell types in scRNA-seq data.

Finally, there are several limitations to the work currently presented in this paper. First, due to issues with parallelisation we could not include the promising recent SMaSH method. Second, we only use the splat method for simulation which does not estimate clustering parameters from real datasets. Third, future work could assess the performance of methods that take into account sample-level replication, such as mixed models and recent pseudobulk methods^[4]^.

## Conclusion

We present a systematic and comprehensive comparison of methods for selecting marker genes in scRNA-seq data. Our results highlight a lack of high concordance between methods, large differences in computational demands and large differences in performance, which occur across simulation, expert-annotated gene and prediction-based analyses. Overall, our results suggest that methods based on logistic regression, Student’s t-test and the Wilcoxon rank-sum test all have strong performance. On the other hand, the scran `findMarkers()`, Cepo and NSForest methods showed generally poor performance across comparisons. Our discovery of unexpected and undocumented behaviour in several cases highlights the importance of continued scrutinising of widely-used methods for scRNA-seq data analysis.

## Methods

### Marker gene selection methods

In this section we provide more details about the marker gene selection methods compared in this paper.

#### Method parameters

##### Seurat

We change the `test.use` parameter that controls which two sample statistical test is performed between the cluster of interest and all other clusters. The parameters currently can take the following values:

- `wilcox:` A Wilcoxon rank-sum test (**Default**);
- `t:` Welch’s t-test;
- `bimod:` Likelihood ratio test for single-cell gene expression^[35]^;
- `DESeq2:` The R package DESeq2 (excluded from comparisons, see below);
- `LR:` A logistic regression is fit for each gene with a binary cluster of interest indicator as the response;
- `MAST:` The R package MAST package^[13]^;
- `negbinom:` A two-group negative binomial GLM;
- `poisson:` A two-group Poisson GLM;
- `roc:` For each gene a classifier is built based on that gene’s ability to discriminate between clusters and evaluated using the AUC. The method ranks genes by predictive power 2|AUC – 0.5|.

Seurat corrects the p-values returned by its methods for multiple testing using Bonferroni’s correction.

##### Scanpy

We change: the `test.use` parameter, which controls which test is performed, `rankby_abs` which controls whether genes are ranked by the absolute value of, or the raw score, and `tie_correct` which controls whether tie correction is performed in the Wilcoxon rank-sum test.

- `t-test:` Welch’s two-sample t-test (**Default**);
- `t-test_over_estimvar:` Welch’s two sample t-test where the size of the *rest* group is set to the size of the cluster of interest;
- `wilcoxon:` A Wilcoxon rank-sum test;
- `logreg:` Gene-wise logistic regression (excluded from comparisons, see below).

The `rankby_abs` and `tie_correct` parameters both take True/False values.

Scanpy corrects the p-values returned by its methods for multiple testing using the Benjamani-Hochberg method^[36]^

##### scran

The *scran* package has two functions for selecting marker genes. We benchamrk both in this manuscript.

###### findMarkers()

The `findMarkers()` function uses a pairwise DE approach to select marker genes.

In the function we alter: the `test.type` parameter, which controls which two-sample statistical test is performed and `pval.type` which controls how scran aggregates p-values. The `test.type` parameters take the following values:

- `t:` Welch’s two-sample t-test (**Default**);
- `wilcox:` Wilcoxon rank-sum test;
- `binom:` The counts are binarised and an exact binomial test is performed.

The `pval.type` parameter takes values:

- `any:` Allows selection of different sized unions of p-values between the individual tests;
- `all:` Combines the p-values using an intersection union test ^[37]^; practically the largest of the p-values is taken as the overall p-value;
- `some:` A Holm-Bonferroni correction is applied to the p-values is applied and then the middle-most p-value is taken as the overall p-value.

###### scoreMarkers()

The `scoreMarkers` function uses pairwise DE statistic calculation to select marker genes the following statistics are calculated in a pairwise fashion:

- Cohen’s d statistic on log fold-change
- AUC
- Log fold-change in detection proportion (the proption of genes that have non-zero expression)

These statistics are summarised across the pairwise comparisons in five ways:

- Mean
- Median
- Maximum
- Minimum
- Minimum rank

Giving 15 different metrics by which to rank genes for marker gene status and ultimately select marker genes.

### Excluded methods

We exclude the following methods from our benchmarking.

#### Scanpy’s Logistic regression method

The implementation was found to not return what is documented, and it is not tested. Seurat’s logistic regression implementation is also included in the benchmarking.

#### SMaSH

Our benchmarking uses a Snakemake workflow which parallelises jobs (primarily running a specific method on a particular dataset) across a High Performance Computing server. Without this parallelisation running the benchmarking would be computationally infeasible. Unfortunately, we found that in this framework we could not get SMaSH to run successfully. We believe this is because SMaSH writes to a file with a fixed filename during model fitting, so when SMaSH is parallelised the multiple jobs try to write and read to the same file simultaneously, causing errors.

#### Seurat’s DESeq2 method

Seurat’s DESeq2 method was excluded was the comparison after persistent errors getting it to run on the PBMC datasets, apparently due to a low number of counts.

#### COMET

COMET ^[38]^ is a marker gene selection method particularly designed to create panels of markers for functional followup. COMET uses the XL-minimal Hyper-Geometric test to rank genes^[39]^. However, COMET’s implementation has problems which make it difficult to benchmark. First, previous (non-neutral) benchmarking has shown that COMET is extremely computationally intensive^[40]^. (Vargo and Gilbert^[40]^ exclude COMET from their comparisons.) Second, COMET’s Python implementation does not allow the processing of datasets ‘in memory’, instead requiring multiple text files as input. This requirement means using COMET is challenging, particularly due to highly parallelised nature of our benchmarking workflow. For these reasons we COMET is not included in this manuscript.

#### Venice

Venice^[41]^ is a marker gene selection method based on the idea of assessing the performance of gene expression measurements as a predictor for cell type. Unfortunately, when we ran the provided code, we experienced an error which appeared to be caused by a problem in the implementation. As there were no examples, tests or code replicating the results of the manuscript we were unable to establish the cause of the problem.

##### Self-implemented methods

In addition to the existing methods we implemented several methods:

- Random: A random sample of genes are selected as marker genes.
- Log fold-change sorted by absolute value: The top genes ranked by the absolute value of the calculated log fold-change are selected as marker genes.
- Log fold-change sorted by raw value: The top genes ranked by the raw value of the calculated log fold-change are selected as marker genes.
- Student’s t-test: A one-vs-rest two sample Student’s t-test is performed between the cluster of interest and all other clusters. The top marker genes ranked by the test’s p-value are selected as marker genes. Note that the t-tests implemented in scran, Scanpy and Seurat are Welch’s t-test.

For the log fold-change methods we implement Seurat’s formula for the log fold-change using the log-normalised counts as input.

##### Other method implementation details

The edgeR, limma and glmGamPoi methods are all generic methods for DE testing, as such they need additional code to be used to select marker genes. In this paper, all these methods are implemented in a one-vs-rest manner for marker genes. This decision was made based on the poor performance of scran’s methods which implement pairwise testing in early version of the benchmarking.

### Computational experiment details

The workflow management tool Snakemake v6.6.1^[42]^ was used to orchestrate all computational experiments described in the paper. All comparisons were run on the University of Melbourne “Spartan” High Performance Computing cluster^[43]^ which uses the Slurm job scheduler. Analyses were performed in R v4.1.0^[44]^. To run methods implemented in Python via R, the reticulate R package^[45]^ and Python v3.8.6^[46]^ were used. To wrangle data we used the dplyr^[47]^, tidyr^[48]^ and purrr^[49]^ packages. General purpose visualisations were created using the R package ggplot2 ^[50]^ and then assembled into figures using the R package patchwork ^[51]^. The plots that show a comparison of the rankings of multiple methods were created using the `rank_rank_plot()` function in the topconfects R package^[52]^. Plots that show the expression of a gene in different clusters were created using the `plotExpression()` function from the scater R package^[53]^.

### Data processing

All the single-cell datasets were processed in a uniform manner prior to analysis, to remove effects of different preprocessing approaches. All datasets were log-normalised and then subset to the top 2,000 highly variable genes (HVGs) using best practices approaches in the Bioconductor ecosystem^[54]^. For the Lawlor, Smart-seq3, Zeisel and pbmc3k dataset expert-annotated marker genes which are not part of the top 2,000 HVGs were also selected.

### Concordance measurement and selected marker gene characteristics details

To assess the concordance of the methods for each cluster and dataset we selected the (at most) top 10 selected marker genes for each method. For each cluster and dataset the proportion of genes shared between the result of each method and all other methods was calculated. This proportion was then averaged over cluster and dataset to produce a matrix encoding the average proportion of shared genes between methods. This matrix was converted a distance matrix using `dist()` function and a hierarchical clustering was calculated with `hclust()`.

To assess the direction of regulation of the method we selected the (at most) top 20 genes from each method and computed the proportion of these genes that were up-regulated. We defined a gene to be up-regulated if it had a log fold-change, or statistic proportional to the log fold-change, greater than 0. Note that the RankCorr, NSForest and Cepo methods do not output either a log fold-change or statistic proportional to the log fold-change. For these methods we calculate a log fold-change (using Seurat’s formula) and add this to output of the method.

To assess the magnitude of the p-values produced by the methods we select the top 40 genes of each method for every cluster in a given dataset and extract the associated multiple testing corrected p-value output by the methods. (Note that different methods do use different multiple testing correction methods.) Those p-values which took values of exactly 0 were excluded. Seurat’s ROC method, NSForest, Cepo, COSG, RankCorr and (self-implemented) log fold-change and random methods were all excluded from this analysis as they do not produce p-values. In the Zhao dataset all top 40 genes for the Seurat (Poisson GLM) method have exactly-zero p-values so the method was excluded from plotting.

To assess the exactly-zero p-values produced by the methods we calculated the number of exactly-zero p-values produced by each method in every cluster in all datasets. To plot the data the we took the median over clusters for each method and dataset. No limit was set on the number of top marker genes taken as it was observed that in some datasets methods could produce hundreds of exactly-zero p-values. Again, methods that do not produce p-values (see above) were excluded from the analysis.

### Simulated performance assessment

Datasets were simulated using the splatter model implemented in the Splatter package^[27]^. The splat model is designed to be a general purpose simulator of scRNA-seq data. In this paper we adapt it to simulate marker genes by designing and implementing a gene-wise marker gene selection and ranking strategy based on the simulation parameters of the model. This method is designed to select and up-rank genes which visually appear to be marker genes and match our holistic view of the characteristics of good marker genes.

#### Simulated marker genes selection strategy

A key component of the splatter model is the set of DE factors for each group and each gene. Let *g* ∈ {1, …, *G*} index genes and *k* ∈ {1, … *K*} index groups. Then *β_gk_* is the DE indicator for the *k*th group and gth gene. In the splat model *β_gk_* encodes the amount of DE the gth gene shows in the *k*th group relative to a background level of expression. In this context *β_g′k′_* = 1 would indicate background expression for gene *g*′ in group *k*′.To assess whether each gene is a marker gene for a particular cluster we use the following original score, *m_g_*. Let *k*′ be the index of the particular cluster of interest, then:

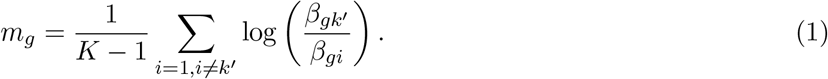

This score is the mean of the log-ratio of the DE factor in the cluster of interest versus all other clusters. In practice the score can consistently up-rank genes that have large log fold-changes. However, if the overall mean of the gene is small the genes do not visually appear to be marker genes. We therefore filter to only genes which show sufficient mean expression: in practice we have found that only keeping genes with gene mean > 0.1 works well.

Our overall strategy for selecting simulated marker genes for a specific cluster from the simulated data consists of:

1. Filter out genes which have simulated mean expression < 0.1;
2. Calculate the marker gene score *m_g_* for each gene;
3. Rank genes by *m_g_*;
4. Select the top *n* genes as marker genes;

for some suitable small *n*. Compared to other methods for selecting marker genes such as the strategy used in the original Splatter paper this score is less restrictive, better accommodating cases where multiple DE factors are not 1 but the gene is still a strong marker gene. The strategy described above can, in theory, be used to select both and up- and down-regulated marker genes. In this paper, however, we only use it to select up-regulated marker genes. While we have many examples of true up-regulated marker genes (all of the expert-annotated maker genes used in this paper are up-regulated) it is not clear what the ideal expression profile of a down-regulated marker gene is. Indeed, when we used the above strategy to select down-regulated marker genes we found that their expression patterns did not visually appear to be marker genes.

#### Simulation parameters

To perform simulations all estimable parameters were estimated from all datasets. To account for the fact that the splat model’s parameter estimation does not account for clustering in the data we filtered each dataset to only one cluster and then performed the parameter estimation on those cells. The cluster used for each dataset is recorded in Table S5. Clusters were mainly chosen based on size, with larger clusters preferred, and to avoid representing an ‘unusual’ cell type. For the three PBMC datasets a B cell cluster was chosen for consistency. Splatter cannot, however, estimate parameters that control the amount of DE between clusters from data. There are two such parameters: the location and scale parameters for the DE factor. In the experiments presented in the main paper we take these parameters to be 3 and 0.2 respectively. These values were chosen as they produced marker genes with similar expression profiles to those observed in real datasets (Figure S7). Additional simulations suggest that the results of method comparison are broadly similar for different values of the location parameter (Figure S9). Note that in all of these analyses we fix the value of the scale parameter; this is because the parameter controls the variance of the (lognormal) distribution and so has less impact on the amount of DE. The dropout and outlier gene probabilities in the splat model were both set to zero. Each simulated dataset was replicated 3 times using a different seed for randomisation.

#### Simulation performance measurement

To measure the performance of the methods we used the following strategy for each cluster in each (replicated) simulated dataset. We selected the (at most) top 20 selected marker genes for each method and the top 20 simulated marker genes. A gene was a true positive if it occurs in both lists, a false positive if it occurs in the list of selected genes but not in the list of simulated genes, a false negative if it does not occur in the selected list of genes but does appear in the simulated list, and a true negative if occurs in neither of the lists. Using these definitions we calculate the number of true positives (TP), false positives (FP) and false negatives (FN) and calculate:

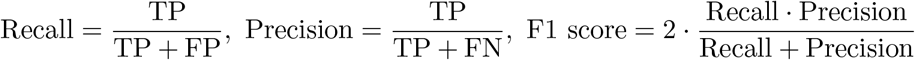

Note that with these definitions the number of simulated genes = TP + FP and the number of selected genes = TP + FN. To summarise the performance of methods for each simulation scenario (i.e. each dataset) we calculated the median recall, precision and F1 score across clusters and simulation replicates.

### Expert-annotated marker gene details

To assess the ability of the methods to recover expert-annotated marker genes we use the following approach. First, we identified sets of expert-annotated marker genes for the Zeisel, Lawlor, pbmc3k and Smart-seq3 datasets:

- *pbmc3k*: The expert-annotated marker genes were taken from the “Seurat - Guided Clustering Tutorial” ^[33]^ (Table S1);
- *Lawlor*: The expert-annotated marker genes were taken from the paper describing the dataset (see the fifth paragraph) ^[32]^. In the paper these genes were used to annotate the defined clustering. (Table S2);
- *Zeisel*: The expert-annotated marker genes were taken from the paper describing the dataset (see the fourth paragraph). We exclude the “None/Other” from expert-annotated marker gene analyses as it does not have marker genes in the original paper.^[55]^. (Table S3);
- *Smart-seq3*: The expert-annotated marker genes were taken from the paper describing the dataset^[56]^. Specifically, these genes were taken from Extended Data Figure 10. Note that not all the marker genes identified in this figure are used. Marker genes for clusters which easily mapped to cluster present in the final version of the clustering were preferred. (Table S4)

Then, to measure the performance of the methods we select marker genes for all clusters in all dataset with expert-annotated marker genes (the number of top genes used was chosen based on the number of expert-annotated marker genes). For each cluster in each dataset the recall of the method was calculated (see above for more information about how this was defined and calculated).

### Predictive performance measurement

To assess the predictive performance of the methods we used the following strategy. For each real dataset we selected the (at most) top 5 marker genes for every cluster, and combined them to give an overall set of genes for each dataset. Either a KNN classifier, with three nearest neighbours, or a SVM classifier, were then trained on only the data from the set of genes for each method, dataset pair. Specifically, we used the `knn` function from the `FNN` R package^[57]^ to implement the KNN classifier and the function `svm` from the `e1071`^[58]^ package to implement the SVM. To assess the performance of the classifier 5-fold cross-validation was used. Each dataset (for each method) was divided into 5 approximately equal groups (i.e. approximately 20% of the total cells) using a sample stratified by cluster. The stratified sample was used to prevent folds having no examples of rare cell types. The model was then fit 5 times with each of the groups being used as the test set while the other groups were pooled to give the training set. Method performance was summarised using the F1 score of the resulting classifier. See above for the mathematical definition of the F1 score. To further summarise the results the median F1 score of the method, dataset pair across clusters was calculated.

### Speed and memory measurement

To measure method speed we used the R `system.time()` function, extracting the elapsed time. To measure the memory usage we used the benchmark functionality from Snakemake, which uses the `ps` command to measure the memory of the submitted jobs. Specifically, we report `max_vss` as the memory. Note that for methods implemented in Python both the time and memory measurements take into account the overhead of converting between R and Python using the reticulate package. The memory measurement also includes various data and output wrangling tasks needed to run specific methods.

### Implementation assessment

To compare the quality of the methods’ implementations we assessed implementation as Good, Adequate or Poor based on five criteria: accessibility (how easy is the software to access i.e. where is it hosted?), installation (how easy is the software to install?), documentation (what is the quality of documentation?), ease of use (how is easy is the software to use for selecting marker genes?) and interpretability of output (how easy is the output to interpret and use?). For further details of how each criteria was rated as Good Adequate or Poor see the Supplementary Methods.

### Log fold-change calculation methods

Let *Y_ig_* be the log-normalised expression value for cell *i* and gene *g*, *G*_1_ and *G*_2_ be the indices of the cells in the cluster of interest and all other clusters respectively, and *n*_1_ and *n*_2_ give the number of cells in the two groups. Then, for a given cluster of interest, Scanpy and Seurat calculate the log-fold change for gene *g* as:

#### Seurat

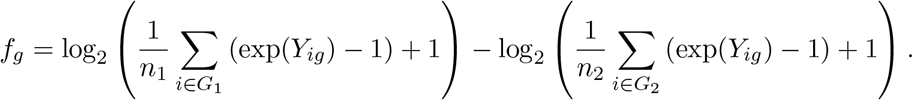

Note that’s Seurat’s Poisson and Negative Binomial GLM methods calculate the log fold-change based on the raw count, that is the above formula applies with *Y_ig_* being the observed count for cell *i* in gene *g*.

#### Scanpy

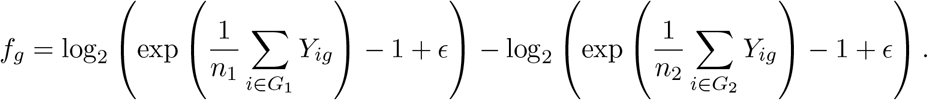

where *ϵ* = 10^−9^. Scanpy’s documentation notes that the log-fold change values are “an approximation calculated from mean-log values” This refers to how the inversion of the log(*x* + 1) transformation of the log-normalised counts by exp(*x*) – 1 is done on the mean of the log-transformed counts, not the log transformed counts themselves. In Scanpy’s implementation the means of the log transformed data are precalculated, so this approximation is probably designed to exploit this feature.

### Relationship between score-based and p-value-based ranking

This equivalence follows from the property that cumulative distribution functions are monotonically increasing. For a two-tailed test of the null hypothesis *H*_0_ : *μ* = 0 which produces a z-score, *z*, the p-value, *p*, is calculated as

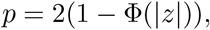

where Φ(·) is the CDF of the standard normal distribution. If there are two scores, *z*_1_ and *z*_2_ say, with |*z*_1_| < |*z*_2_|, then from the fact that Φ(·) is monotonically increasing it can be derived that:

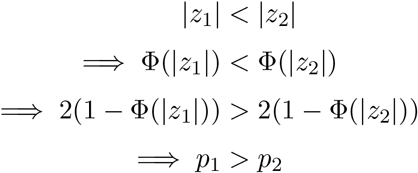

where the first inequality follows from the increasing monotonic property of the CDF. This argument shows that ranking by the absolute value of the z-score (in decreasing order) will be the same as ranking by the p-values (in increasing order) for tests that return a z-score.

This equivalence does not hold, however, for the Welch’s t-test (which is the default statistical test used by Scanpy). In Welch’s t-test the degrees of freedom (*ν*) of the t-distribution that the t-value is compared is dependent on the variance of the data. Different genes will therefore have different *ν* values so each p-value is calculated from a different instance of the t-distribution and the mathematical argument presented for ranking by z-score does not hold.

### Differential expression testing strategy

Different methods based on DE testing use different strategies to perform the testing. When DE testing is used to identify marker genes we need to assess which genes are different in the cluster of interest (i.e. the cluster we wish to select marker genes for) relative to the other clusters present in the data. The *one-vs-rest* and *pairwise* testing strategies are different approaches for quantifying this difference.

#### one-vs-rest

In the one-vs-rest approach the cells in all other clusters and pooled and a statistical test is performed between the cluster of interest and the pooled ‘rest’ cluster for each gene. This method is implemented by Scanpy and Seurat and we the approach we use this approach to implement edgeR, limma and glmGamPoi for marker gene selection in this paper.

#### pairwise

In the pairwise approach a statistical test is performed between the cluster of interest and each other cluster for each gene. This gives *K* – 1 p-values (or statistics) for each gene, cluster of interest combination, where *K* is the number of clusters in the dataset. To get an overall measure of significance for each gene the p-values (or statistics) need to be combined in some way. This method is implemented in scran. The scran `findMarkers()` function performs pairwise DE testing, while the `scoreMarkers()` function calculates a statistic measuring DE in pairwise fashion. The rationale for this more complex DE approach is that the multiple p-values provide more information and that it is hypothetically less affected by the relative sizes of different clusters^[59]^.

### Availability

All code used in the analyses presented in this paper is available at https://gitlab.svi.edu.au/biocellgen-public/mage_2020_marker-gene-benchmarking. The raw datasets were accessed from the following sources. The Lawlor, Zhao and Zeisel datasets were accessed from the `scRNAseq` R package^[60]^. The Paul dataset was accessed from the Scanpy package. The Astrocyte, Smart-seq3 and CITE-seq datasets were accessed from the data made available with the associated publication. The pbmc3k dataset was accessed from the `TENxPBMCData` R package^[61]^. All processed datasets (except the Endothelial and Mesenchymal datasets which are restricted pending their formal publication) are available through Zenodo: doi:10.5281/zenodo.6513335.

## Author contributions

JMP conceived the project. JMP designed the comparisons and simulations, implemented the benchmarking and performed analysis, supervised by DJM. JMP and DJM wrote the manuscript.

## Acknowledgements

The authors would like to thank Nicholas Banovich and Jonathan Kropski for providing the unpublished Endothelial and Mesenchymal datasets and for helpful comments on the manuscript, Christina Azodi for many helpful discussions, Sagrika Chugh for advice concerning the Endothelial dataset, and all other members of the Bioinformatics and Cellular Genomics Group at the St Vincent’s Institute of Medical Research and the University of Melbourne for their comments and support. We acknowledge and thank the patients who contributed samples for research that make up the datasets used in this paper.

## Tables

## References

[1] Valentine Svensson, Eduardo da Veiga Beltrame, and Lior Pachter. A curated database reveals trends in single-cell transcriptomics. Database, 2020(baaa073), January 2020. ISSN 1758-0463. doi:10.1093/database/baaa073.

[2] Luke Zappia, Belinda Phipson, and Alicia Oshlack. Exploring the single-cell RNA-seq analysis landscape with the scRNA-tools database. PLOS Computational Biology, 14(6):e1006245, June 2018. ISSN 1553-7358. doi:10.1371/journal.pcbi.1006245.

[3] Luke Zappia and Fabian J. Theis. Over 1000 tools reveal trends in the single-cell RNA-seq analysis landscape. Genome Biology, 22(1):301, October 2021. ISSN 1474-760X. doi:10.1186/s13059-021-02519-4.

[4] Helena L. Crowell, Charlotte Soneson, Pierre-Luc Germain, Daniela Calini, Ludovic Collin, Catarina Raposo, Dheeraj Malhotra, and Mark D. Robinson. Muscat detects subpopulation-specific state transitions from multi-sample multi-condition single-cell transcriptomics data. Nature Communications, 11(1):6077, November 2020. ISSN 2041-1723. doi:10.1038/s41467-020-19894-4.

[5] MGP van der Wijst, DH de Vries, HE Groot, G Trynka, CC Hon, MJ Bonder, O Stegle, MC Nawijn, Y Idaghdour, P van der Harst, CJ Ye, J Powell, FJ Theis, A Mahfouz, M Heinig, and L Franke. The single-cell eQTLGen consortium. eLife, 9:e52155, March 2020. ISSN 2050-084X. doi:10.7554/eLife.52155.

[6] Gabriela Balderrama-Gutierrez, Heidi Liang, Narges Rezaie, Klebea Carvalho, Stefania Forner, Dina Mattheos, Elisabeth Rebboah, Kim N. Green, Andrea J. Tenner, Frank LaFerla, and Ali Mortazavi. Single-cell and nucleus RNA-seq in a mouse model of AD reveal activation of distinct glial subpopulations in the presence of plaques and tangles, October 2021.

[7] M. E. Nelson, S. G. Riva, and A. Cvejic. SMaSH: A scalable, general marker gene identification framework for single-cell RNA-sequencing. BMC Bioinformatics, 23(1):328, August 2022. ISSN 1471-2105. doi:10.1186/s12859-022-04860-2.

[8] Vitalii Kleshchevnikov, Artem Shmatko, Emma Dann, Alexander Aivazidis, Hamish W. King, Tong Li, Artem Lomakin, Veronika Kedlian, Mika Sarkin Jain, Jun Sung Park, Lauma Ramona, Elizabeth Tuck, Anna Arutyunyan, Roser Vento-Tormo, Moritz Gerstung, Louisa James, Oliver Stegle, and Omer Ali Bayraktar. Comprehensive mapping of tissue cell architecture via integrated single cell and spatial transcriptomics, November 2020.

[9] Hannah A. Pliner, Jay Shendure, and Cole Trapnell. Supervised classification enables rapid annotation of cell atlases. Nature Methods, 16(10):983–986, October 2019. ISSN 1548-7105. doi:10.1038/s41592-019-0535-3.

[10] Ze Zhang, Danni Luo, Xue Zhong, Jin Huk Choi, Yuanqing Ma, Stacy Wang, Elena Mahrt, Wei Guo, Eric W Stawiski, Zora Modrusan, Somasekar Seshagiri, Payal Kapur, Gary C. Hon, James Brugarolas, and Tao Wang. SCINA: A Semi-Supervised Subtyping Algorithm of Single Cells and Bulk Samples. Genes, 10(7):531, July 2019. ISSN 2073-4425. doi:10.3390/genes10070531.

[11] Hongyu Guo and Jun Li. scSorter: Assigning cells to known cell types according to marker genes. Genome Biology, 22(1):69, February 2021. ISSN 1474-760X. doi:10.1186/s13059-021-02281-7.

[12] Charlotte Soneson and Mark D. Robinson. Bias, robustness and scalability in single-cell differential expression analysis. Nature Methods, 15(4):255–261, April 2018. ISSN 1548-7105. doi:10.1038/nmeth.4612.

[13] Greg Finak, Andrew McDavid, Masanao Yajima, Jingyuan Deng, Vivian Gersuk, Alex K. Shalek, Chloe K. Slichter, Hannah W. Miller, M. Juliana McElrath, Martin Prlic, Peter S. Linsley, and Raphael Gottardo. MAST: A flexible statistical framework for assessing transcriptional changes and characterizing heterogeneity in single-cell RNA sequencing data. Genome Biology, 16(1):278, December 2015. ISSN 1474-760X. doi:10.1186/s13059-015-0844-5.

[14] Jordan W. Squair, Matthieu Gautier, Claudia Kathe, Mark A. Anderson, Nicholas D. James, Thomas H. Hutson, Rémi Hudelle, Taha Qaiser, Kaya J. E. Matson, Quentin Barraud, Ariel J. Levine, Gioele La Manno, Michael A. Skinnider, and Grégoire Courtine. Confronting false discoveries in single-cell differential expression. Nature Communications, 12(1):5692, September 2021. ISSN 2041-1723. doi:10.1038/s41467-021-25960-2.

[15] Kip D. Zimmerman, Mark A. Espeland, and Carl D. Langefeld. A practical solution to pseudoreplication bias in single-cell studies. Nature Communications, 12(1):738, February 2021. ISSN 2041-1723. doi:10.1038/s41467-021-21038-1.

[16] Colin Megill, Bruce Martin, Charlotte Weaver, Sidney Bell, Lia Prins, Seve Badajoz, Brian McCandless, Angela Oliveira Pisco, Marcus Kinsella, Fiona Griffin, Justin Kiggins, Genevieve Haliburton, Arathi Mani, Matthew Weiden, Madison Dunitz, Maximilian Lombardo, Timmy Huang, Trent Smith, Signe Chambers, Jeremy Freeman, Jonah Cool, and Ambrose Carr. Cellxgene: A performant, scalable exploration platform for high dimensional sparse matrices, April 2021.

[17] Tianyu Wang, Boyang Li, Craig E. Nelson, and Sheida Nabavi. Comparative analysis of differential gene expression analysis tools for single-cell RNA sequencing data. BMC Bioinformatics, 20(1):40, January 2019. ISSN 1471-2105. doi:10.1186/s12859-019-2599-6.

[18] Yuhan Hao, Stephanie Hao, Erica Andersen-Nissen, William M. Mauck, Shiwei Zheng, Andrew Butler, Maddie J. Lee, Aaron J. Wilk, Charlotte Darby, Michael Zager, Paul Hoffman, Marlon Stoeckius, Efthymia Papalexi, Eleni P. Mimitou, Jaison Jain, Avi Srivastava, Tim Stuart, Lamar M. Fleming, Bertrand Yeung, Angela J. Rogers, Juliana M. McElrath, Catherine A. Blish, Raphael Gottardo, Peter Smibert, and Rahul Satija. Integrated analysis of multimodal single-cell data. Cell, 184(13):3573–3587.e29, June 2021. ISSN 0092-8674, 1097-4172. doi:10.1016/j.cell.2021.04.048.

[19] F. Alexander Wolf, Philipp Angerer, and Fabian J. Theis. SCANPY: Large-scale single-cell gene expression data analysis. Genome Biology, 19(1):15, February 2018. ISSN 1474-760X. doi:10.1186/s13059-017-1382-0.

[20] Brian D. Aevermann, Yun Zhang, Mark Novotny, Mohamed Keshk, Trygve E. Bakken, Jeremy A. Miller, Rebecca D. Hodge, Boudewijn Lelieveldt, Ed S. Lein, and Richard H. Scheuermann. A machine learning method for the discovery of minimum marker gene combinations for cell-type identification from single-cell RNA sequencing. Genome Research, page gr.275569.121, June 2021. ISSN 1088-9051, 1549-5469. doi:10.1101/gr.275569.121.

[21] Min Dai, Xiaobing Pei, and Xiu-Jie Wang. Accurate and fast cell marker gene identification with COSG. Briefings in Bioinformatics, 23(2):bbab579, March 2022. ISSN 1477-4054. doi:10.1093/bib/bbab579.

[22] Michael B. Cole, Davide Risso, Allon Wagner, David DeTomaso, John Ngai, Elizabeth Purdom, Sandrine Dudoit, and Nir Yosef. Performance Assessment and Selection of Normalization Procedures for Single-Cell RNA-Seq. Cell Systems, 8(4):315–328.e8, April 2019. ISSN 2405-4712. doi:10.1016/j.cels.2019.03.010.

[23] Wouter Saelens, Robrecht Cannoodt, Helena Todorov, and Yvan Saeys. A comparison of single-cell trajectory inference methods. Nature Biotechnology, 37(5):547–554, May 2019. ISSN 1546-1696. doi:10.1038/s41587-019-0071-9.

[24] Zhanying Feng, Xianwen Ren, Yuan Fang, Yining Yin, Chutian Huang, Yimin Zhao, and Yong Wang. scTIM: Seeking cell-type-indicative marker from single cell RNA-seq data by consensus optimization. Bioinformatics, 36(8):2474–2485, April 2020. ISSN 1367-4803. doi:10.1093/bioinformatics/btz936.

[25] Bianca Dumitrascu, Soledad Villar, Dustin G. Mixon, and Barbara E. Engelhardt. Optimal marker gene selection for cell type discrimination in single cell analyses. Nature Communications, 12(1):1186, February 2021. ISSN 2041-1723. doi:10.1038/s41467-021-21453-4.

[26] Alexander H. S. Vargo and Anna C. Gilbert. A rank-based marker selection method for high throughput scRNA-seq data. BMC Bioinformatics, 21(1):477, October 2020. ISSN 1471-2105. doi:10.1186/s12859-020-03641-z.

[27] Luke Zappia, Belinda Phipson, and Alicia Oshlack. Splatter: Simulation of single-cell RNA sequencing data. Genome Biology, 18(1):174, September 2017. ISSN 1474-760X. doi:10.1186/s13059-017-1305-0.

[28] Helena L. Crowell, Sarah X. Morillo Leonardo, Charlotte Soneson, and Mark D. Robinson. Built on sand: The shaky foundations of simulating single-cell RNA sequencing data, February 2022.

[29] Junyue Cao, Malte Spielmann, Xiaojie Qiu, Xingfan Huang, Daniel M. Ibrahim, Andrew J. Hill, Fan Zhang, Stefan Mundlos, Lena Christiansen, Frank J. Steemers, Cole Trapnell, and Jay Shendure. The single-cell transcriptional landscape of mammalian organogenesis. Nature, 566(7745):496–502, February 2019. ISSN 1476-4687. doi:10.1038/s41586-019-0969-x.

[30] Jesse M. Zhang, Govinda M. Kamath, and David N. Tse. Valid Post-clustering Differential Analysis for Single-Cell RNA-Seq. Cell Systems, 9(4):383–392.e6, October 2019. ISSN 2405-4712. doi:10.1016/j.cels.2019.07.012.

[31] Oscar Franzén, Li-Ming Gan, and Johan L M Björkegren. PanglaoDB: A web server for exploration of mouse and human single-cell RNA sequencing data. Database, 2019(baz046), January 2019. ISSN 1758-0463. doi:10.1093/database/baz046.

[32] Nathan Lawlor, Joshy George, Mohan Bolisetty, Romy Kursawe, Lili Sun, V. Sivakamasundari, Ina Kycia, Paul Robson, and Michael L. Stitzel. Single-cell transcriptomes identify human islet cell signatures and reveal cell-type–specific expression changes in type 2 diabetes. Genome Research, November 2016. ISSN 1088-9051, 1549-5469. doi:10.1101/gr.212720.116.

[33] Seurat-Guided Clustering Tutorial. https://satijalab.org/seurat/articles/pbmc3k_tutorial.html.

[34] Lucy L. Gao, Jacob Bien, and Daniela Witten. Selective Inference for Hierarchical Clustering. arXiv:2012.02936 [stat], December 2020.

[35] Andrew McDavid, Greg Finak, Pratip K. Chattopadyay, Maria Dominguez, Laurie Lamoreaux, Steven S. Ma, Mario Roederer, and Raphael Gottardo. Data exploration, quality control and testing in single-cell qPCR-based gene expression experiments. Bioinformatics, 29(4):461–467, February 2013. ISSN 1367-4803. doi:10.1093/bioinformatics/bts714.

[36] Yoav Benjamini and Yosef Hochberg. Controlling the False Discovery Rate: A Practical and Powerful Approach to Multiple Testing. Journal of the Royal Statistical Society. Series B (Methodological), 57(1):289–300, 1995. ISSN 0035-9246.

[37] Roger L. Berger and Jason C. Hsu. Bioequivalence trials, intersection-union tests and equivalence confidence sets. Statistical Science, 11(4):283–319, November 1996. ISSN 0883-4237, 2168-8745. doi:10.1214/ss/1032280304.

[38] Conor Delaney, Alexandra Schnell, Louis V Cammarata, Aaron Yao-Smith, Aviv Regev, Vijay K Kuchroo, and Meromit Singer. Combinatorial prediction of marker panels from single-cell transcriptomic data. Molecular Systems Biology, 15(10):e9005, October 2019. ISSN 1744-4292. doi:10.15252/msb.20199005.

[39] Florian Wagner. The XL-mHG Test For Enrichment: A Technical Report, September 2015.

[40] Alexander H. S. Vargo and Anna C. Gilbert. A rank-based marker selection method for high throughput scRNA-seq data. BMC Bioinformatics, 21(1):477, October 2020. ISSN 1471-2105. doi:10.1186/s12859-020-03641-z.

[41] Hy Vuong, Thao Truong, Tan Phan, and Son Pham. Venice: A New Algorithm for Finding Marker Genes in Single-Cell Transcriptomic Data. bioRxiv, page 2020.11.16.384479, November 2020. doi:10.1101/2020.11.16.384479.

[42] Johannes Köster and Sven Rahmann. Snakemake—a scalable bioinformatics workflow engine. Bioinformatics, 28(19):2520–2522, October 2012. ISSN 1367-4803. doi:10.1093/bioinformatics/bts480.

[43] Spartan HPC-Cloud Hybrid: Delivering Performance and Flexibility. https://melbourne.figshare.com/articles/online_resource/Spartan_HPC-Cloud_Hybrid_Delivering_Performance_and_Flexibility/4768291/1, April 2017.

[44] R Core Team. R: A Language and Environment for Statistical Computing. R Foundation for Statistical Computing, Vienna, Austria, 2021. URL https://www.R-project.org/.

[45] Kevin Ushey, JJ Allaire, and Yuan Tang. reticulate: Interface to ‘Python’, 2021. URL https://CRAN.R-project.org/package=reticulate. R package version 1.20.

[46] Guido Van Rossum and Fred L Drake Jr. Python tutorial. Centrum voor Wiskunde en Informatica Amsterdam, The Netherlands, 1995.

[47] Hadley Wickham, Romain François, Lionel Henry, and Kirill Müller. dplyr: A Grammar of Data Manipulation, 2021. URL https://CRAN.R-project.org/package=dplyr. R package version 1.0.7.

[48] Hadley Wickham. tidyr: Tidy Messy Data, 2021. URL https://CRAN.R-project.org/package=tidyr. R package version 1.1.4.

[49] Lionel Henry and Hadley Wickham. purrr: Functional Programming Tools, 2020. URL https://CRAN.R-project.org/package=purrr. R package version 0.3.4.

[50] Hadley Wickham. ggplot2: Elegant Graphics for Data Analysis. Springer-Verlag New York, 2016. ISBN 978-3-319-24277-4. URL https://ggplot2.tidyverse.org.

[51] Thomas Lin Pedersen. patchwork: The Composer of Plots, 2020. URL https://CRAN.R-project.org/package=patchwork. R package version 1.1.1.

[52] Paul F. Harrison, Andrew D. Pattison, David R. Powell, and Traude H. Beilharz. Topconfects: A package for confident effect sizes in differential expression analysis provides a more biologically useful ranked gene list. Genome Biology, 20(1):67, March 2019. ISSN 1474-760X. doi:10.1186/s13059-019-1674-7.

[53] Davis J. McCarthy, Kieran R. Campbell, Aaron T. L. Lun, and Quin F. Wills. Scater: Pre-processing, quality control, normalization and visualization of single-cell RNA-seq data in R. Bioinformatics, 33(8):1179–1186, April 2017. ISSN 1367-4803. doi:10.1093/bioinformatics/btw777.

[54] Aaron T.L. Lun, Davis J. McCarthy, and John C. Marioni. A step-by-step workflow for low-level analysis of single-cell RNA-seq data with Bioconductor. F1000Research, 5:2122, October 2016. ISSN 2046-1402. doi:10.12688/f1000research.9501.2.

[55] Amit Zeisel, Ana B. Muñoz-Manchado, Simone Codeluppi, Peter Lönnerberg, Gioele La Manno, Anna Juréus, Sueli Marques, Hermany Munguba, Liqun He, Christer Betsholtz, Charlotte Rolny, Gonçalo Castelo-Branco, Jens Hjerling-Leffler, and Sten Linnarsson. Brain structure. Cell types in the mouse cortex and hippocampus revealed by single-cell RNA-seq. Science (New York, N.Y.), 347(6226):1138–1142, March 2015. ISSN 1095-9203. doi:10.1126/science.aaa1934.

[56] Michael Hagemann-Jensen, Christoph Ziegenhain, Ping Chen, Daniel Ramsköld, Gert-Jan Hendriks, Anton J. M. Larsson, Omid R. Faridani, and Rickard Sandberg. Single-cell RNA counting at allele and isoform resolution using Smart-seq3. Nature Biotechnology, 38(6):708–714, June 2020. ISSN 1546-1696. doi:10.1038/s41587-020-0497-0.

[57] Alina Beygelzimer, Sham Kakadet, John Langford, Sunil Arya, David Mount, and Shengqiao Li. FNN: Fast Nearest Neighbor Search Algorithms and Applications, 2019. URL https://CRAN.R-project.org/package=FNN. R package version 1.1.3.

[58] David Meyer, Evgenia Dimitriadou, Kurt Hornik, Andreas Weingessel, and Friedrich Leisch. e1071: Misc Functions of the Department of Statistics, Probability Theory Group (Formerly: E1071), TU Wien, 2022. URL https://CRAN.R-project.org/package=e1071. R package version 1.7-11.

[59] Robert A. Amezquita, Aaron T. L. Lun, Etienne Becht, Vince J. Carey, Lindsay N. Carpp, Ludwig Geistlinger, Federico Marini, Kevin Rue-Albrecht, Davide Risso, Charlotte Soneson, Levi Waldron, Hervé Pagès, Mike L. Smith, Wolfgang Huber, Martin Morgan, Raphael Gottardo, and Stephanie C. Hicks. Orchestrating single-cell analysis with Bioconductor. Nature Methods, 17(2):137–145, February 2020. ISSN 1548-7105. doi:10.1038/s41592-019-0654-x.

[60] Davide Risso and Michael Cole. scRNAseq: Collection of Public Single-Cell RNA-Seq Datasets, 2021. R package version 2.6.1.

[61] Kasper D. Hansen, Davide Risso, and Stephanie Hicks. TENxPBMCData: PBMC data from 10X Genomics, 2021. R package version 1.10.0.

[62] Steven P. Lund, Dan Nettleton, Davis J. McCarthy, and Gordon K. Smyth. Detecting Differential Expression in RNA-sequence Data Using Quasi-likelihood with Shrunken Dispersion Estimates. Statistical Applications in Genetics and Molecular Biology, 11(5), October 2012. ISSN 10.1515/1544-6115.1826. doi:10.1515/1544-6115.1826.

[63] Davis J. McCarthy, Yunshun Chen, and Gordon K. Smyth. Differential expression analysis of multifactor RNA-Seq experiments with respect to biological variation. Nucleic Acids Research, 40(10):4288–4297, May 2012. ISSN 0305-1048. doi:10.1093/nar/gks042.

[64] Constantin Ahlmann-Eltze and Wolfgang Huber. glmGamPoi: Fitting Gamma-Poisson generalized linear models on single cell count data. Bioinformatics, 36(24):5701–5702, December 2020. ISSN 1367-4803. doi:10.1093/bioinformatics/btaa1009.

[65] Gordon K. Smyth. Linear Models and Empirical Bayes Methods for Assessing Differential Expression in Microarray Experiments. Statistical Applications in Genetics and Molecular Biology, 3(1), February 2004. ISSN 10.2202/1544-6115.1027. doi:10.2202/1544-6115.1027.

[66] Charity W. Law, Yunshun Chen, Wei Shi, and Gordon K. Smyth. Voom: Precision weights unlock linear model analysis tools for RNA-seq read counts. Genome Biology, 15(2):R29, February 2014. ISSN 1474-760X. doi:10.1186/gb-2014-15-2-r29.

[67] Hani Jieun Kim, Kevin Wang, Carissa Chen, Yingxin Lin, Patrick P. L. Tam, David M. Lin, Jean Y. H. Yang, and Pengyi Yang. Uncovering cell identity through differential stability with Cepo. Nature Computational Science, 1(12):784–790, December 2021. ISSN 2662-8457. doi:10.1038/s43588-021-00172-2.

[68] Franziska Paul, Ya’ara Arkin, Amir Giladi, Diego Adhemar Jaitin, Ephraim Kenigsberg, Hadas Keren-Shaul, Deborah Winter, David Lara-Astiaso, Meital Gury, Assaf Weiner, Eyal David, Nadav Cohen, Felicia Kathrine Bratt Lauridsen, Simon Haas, Andreas Schlitzer, Alexander Mildner, Florent Ginhoux, Steffen Jung, Andreas Trumpp, Bo Torben Porse, Amos Tanay, and Ido Amit. Transcriptional Heterogeneity and Lineage Commitment in Myeloid Progenitors. Cell, 163(7):1663–1677, December 2015. ISSN 0092-8674, 1097-4172. doi:10.1016/j.cell.2015.11.013.

[69] Jun Zhao, Ariel Jaffe, Henry Li, Ofir Lindenbaum, Esen Sefik, Ruaidhrí Jackson, Xiuyuan Cheng, Richard A. Flavell, and Yuval Kluger. Detection of differentially abundant cell subpopulations in scRNA-seq data. Proceedings of the National Academy of Sciences, 118(22), June 2021. ISSN 0027-8424, 1091–6490. doi:10.1073/pnas.2100293118.

[70] Marlon Stoeckius, Christoph Hafemeister, William Stephenson, Brian Houck-Loomis, Pratip K. Chattopadhyay, Harold Swerdlow, Rahul Satija, and Peter Smibert. Simultaneous epitope and transcriptome measurement in single cells. Nature Methods, 14(9):865–868, September 2017. ISSN 1548-7105. doi:10.1038/nmeth.4380.

