## Supplementary tables and figures for "A comparison of marker gene selection methods for single-cell RNA sequencing data"

September 13, 2022

### Supplementary methods

#### Implementation criteria

The criteria used to rate the implementations of the methods.

- Accessibility.
  - Good: The software is hosted in standard repository (e.g. Bioconductor, pip)
  - Adequate: The software is only hosted in a standard Git server
  - Poor: The software is hosted in a non-standard way or is not accessible
- Installation. How easy is the software to install?
  - Good: The software can be installed in a standard way.
  - Adequate: The software can be installed in a non-standard way.
  - Poor: The software can only be installed with difficulty or not at all.
- Documentation quality. How well documented is the software?
  - Good: There is substantial documentation
  - Adequate: There is some documentation; enough to run the software.
  - Poor: There is little or no documentation.
- Ease of use. How easy is it to run the software to select marker genes?
  - Good: The package natively supports selecting marker genes and common scRNA-seq data formats are supported.
  - Adequate: The package natively supports selecting marker genes but the passed data is is required in an odd format.
  - Poor: Additional code must be written to support the selection of marker genes
- Quality of output. How easy is it to use and interpret the output?
  - Good: The output is in a sensible format with all expect components
  - Adequate: The output is in a mostly sensible format, possibly with some components mangled.
  - Poor: The output is in an odd, difficult to use format.

### Supplementary figures and tables

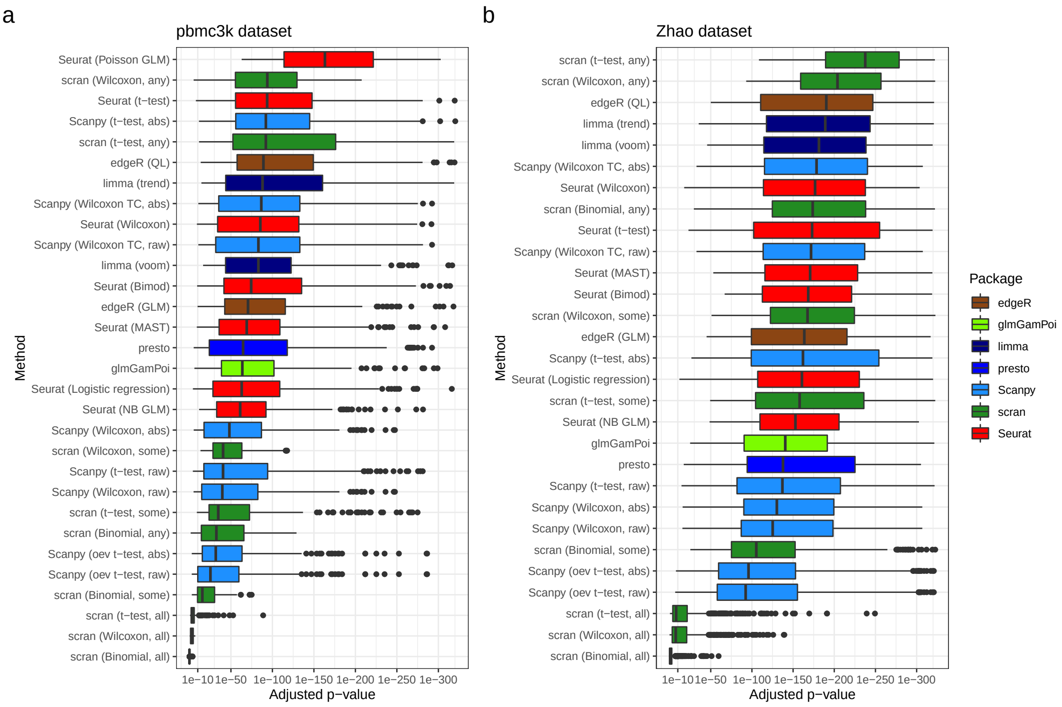


**Figure S1: Adjusted p-value magnitudes**. a) Boxplots of (multiple correction) adjusted p-values for all methods which output p-values run on the pbmc3k dataset (results for all clusters included in each boxplot). b) As in a) for the Zhao dataset. Different methods use different methods to perform multiple testing correction. (See Methods).

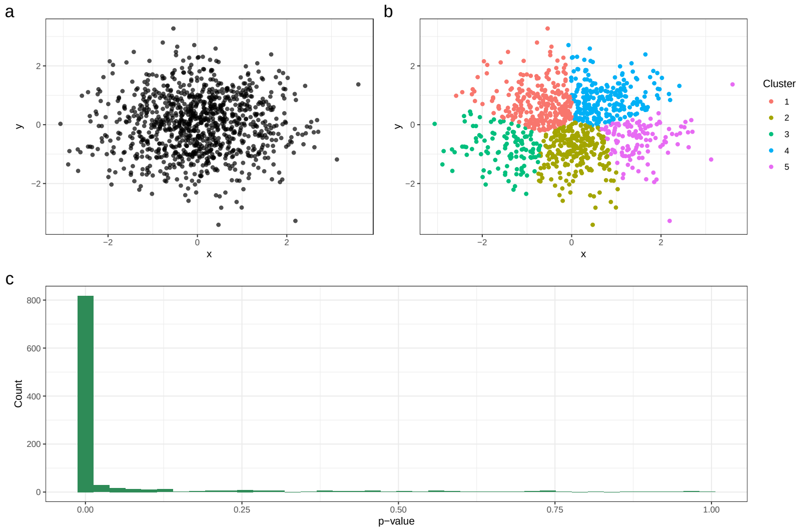


**Figure S2: A graphical illustration of the problem of ‘double dipping’ - performing inference after clustering**. a) 1000 points simulated from a bivariate normal distribution with means of zero and identity covariance matrix. b) A $k$-means clustering with $k = 5$ applied to the dataset in a). c): Histogram of 1000 p-values computed by performing Welch’s t-test between the $y$ values of cluster 1 and the $y$ values of all other clusters in 1000 simulated replicates (same analysis) of the dataset from b). Despite no different in the generative models creating the data in cluster 1 and all other clusters, the computed p-values are generally highly significant.

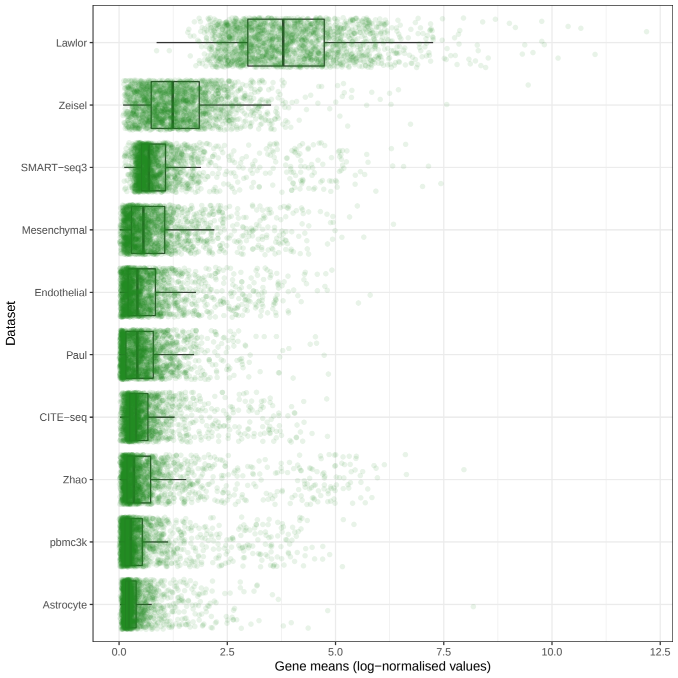


**Figure S3: Gene means across datasets**. The distribution of gene means across datasets calculated using on log-normalised counts. The Lawlor dataset has substantially higher mean expression compared to other datasets.

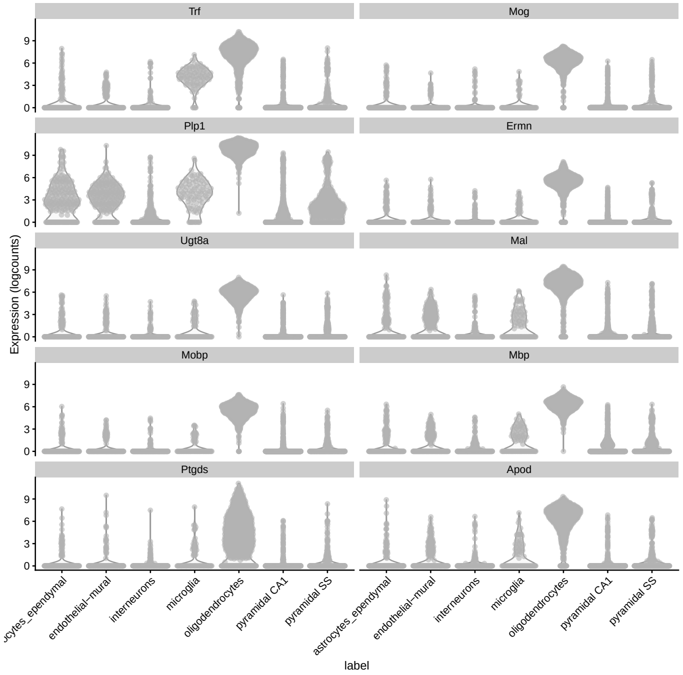


**Figure S4: Ten top marker genes selected by the Seurat (Wilcoxon) method in the Oligodendrocyte cluster, Zeisel dataset**

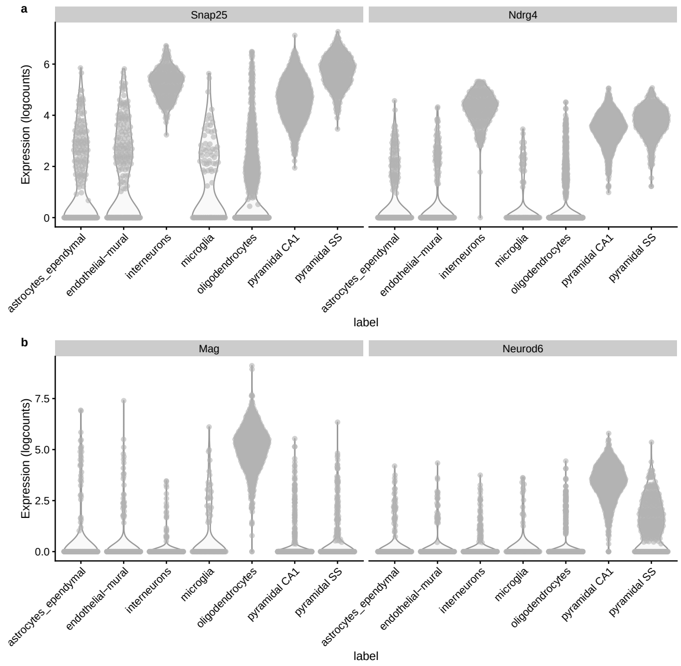


**Figure S5: Top two marker genes selected by the Seurat (t-test) and Scran t-test-any method in the Interneuron cluster, Zeisel dataset** a) Seurat t-test selected marker genes. b) Scran t-test any selected marker genes.

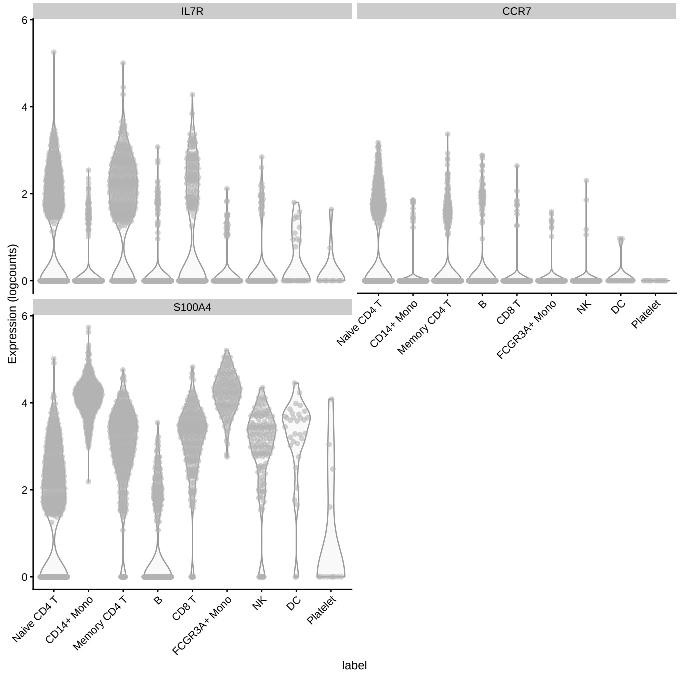


**Figure S6: Expert annotated marker genes for the CD4 T Memory and CD4 T Naive cells in the pbmc3k datasets** Note that *IL7R* is annotated as a marker gene for both clusters

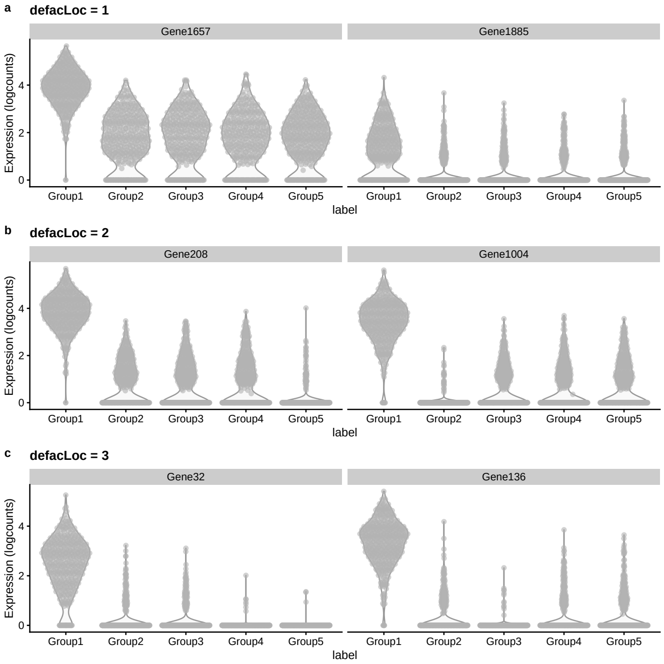


**Figure S7: Top 2 'true' marker genes for simulation with different values of** `de.facLoc` All simulations have all other parameters estimated from the pbmc3k dataset and have `de.facScale` = 0.2. The `de.facScale` parameter controls the variance of the distribution of DE factors and therefore has less of an effect on the simulations.

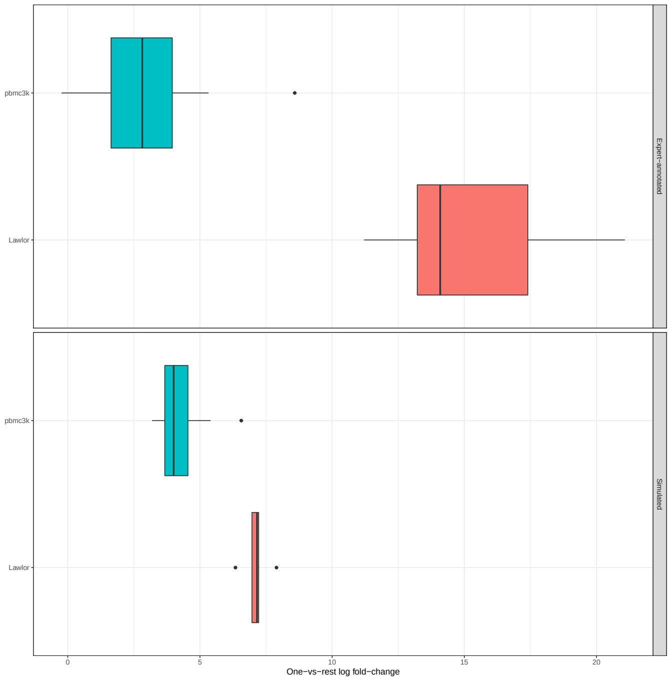


**Figure S8: Log fold-change values for the top simulated and expert-annotated genes in the pbmc3k and Lawlor datasets**. The simulated and expert-annotated marker genes have similar log fold-change values, and the simulations are able to partially recapitulate the differing magnitude of the log fold-changes in the two datasets. To make the number of simulated and expert-annotated marker genes similar, the top 3 simulated marker genes for the pbmc3k dataset and the top simulated marker genes for the Lawlor dataset are used.

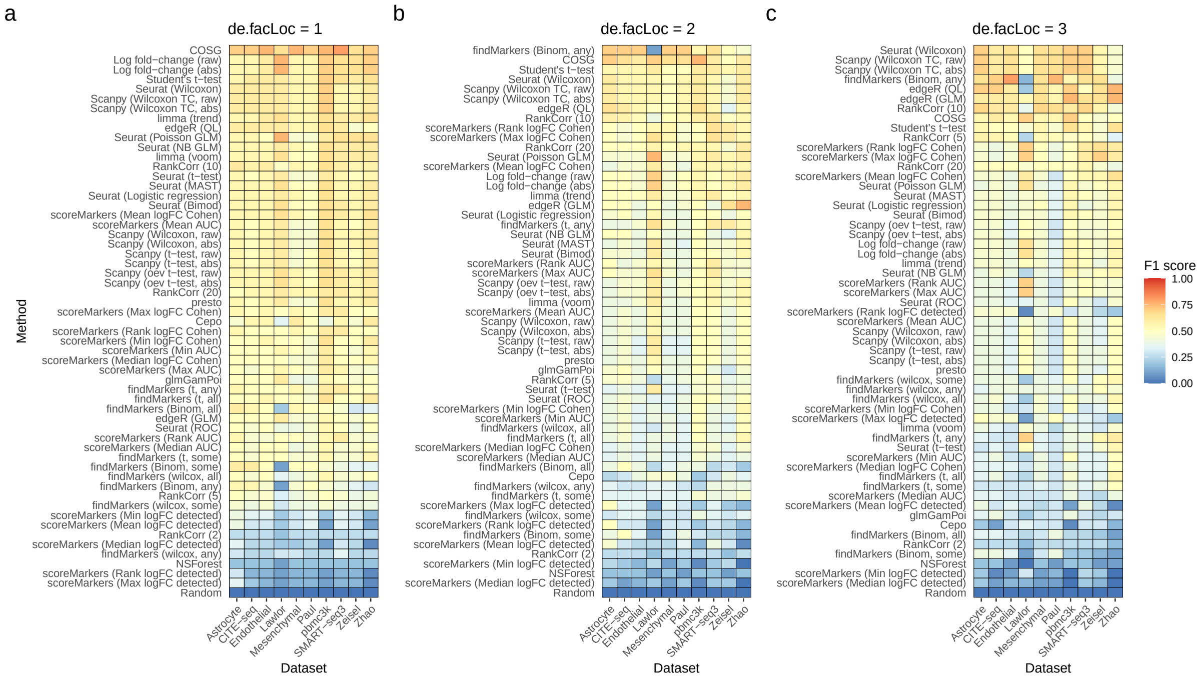


**Figure S9: Simulation results using different values of splatter de.facLoc parameters** Results are relatively concordant for different values of the parameter. In a) when `de.facLoc` = 1 scran Binomial-any shows poor performance and the methods ranking by log fold-change show much better results. All simulations have `de.facScale` = 0.2

**Table S1:** Expert marker genes: pbmc3k

| Cell type | Marker genes |
|---|---|
| Naive CD4+ T | IL7R, CCR7 |
| CD14+ Mono | CD14, LYZ |
| Memory CD4+ | IL7R, S100A4 |
| B | MS4A1 |
| CD8+ T | CD8A |
| FCGR3A+ Mono | FCGR3A, MS4A7 |
| NK | GNLY, NKG7 |
| DC | FCER1A, CST3 |
| Platelet | PPBP |

**Table S2:** Expert marker genes: Lawlor

| Cell type | Marker gene |
|---|---|
| Beta | INS |
| Stellate | COL1A1 |
| Ductal | KRT19 |
| Alpha | GCG |
| Acinar | PRSS1 |
| Gamma/PP | PPY |
| Delta | SST |

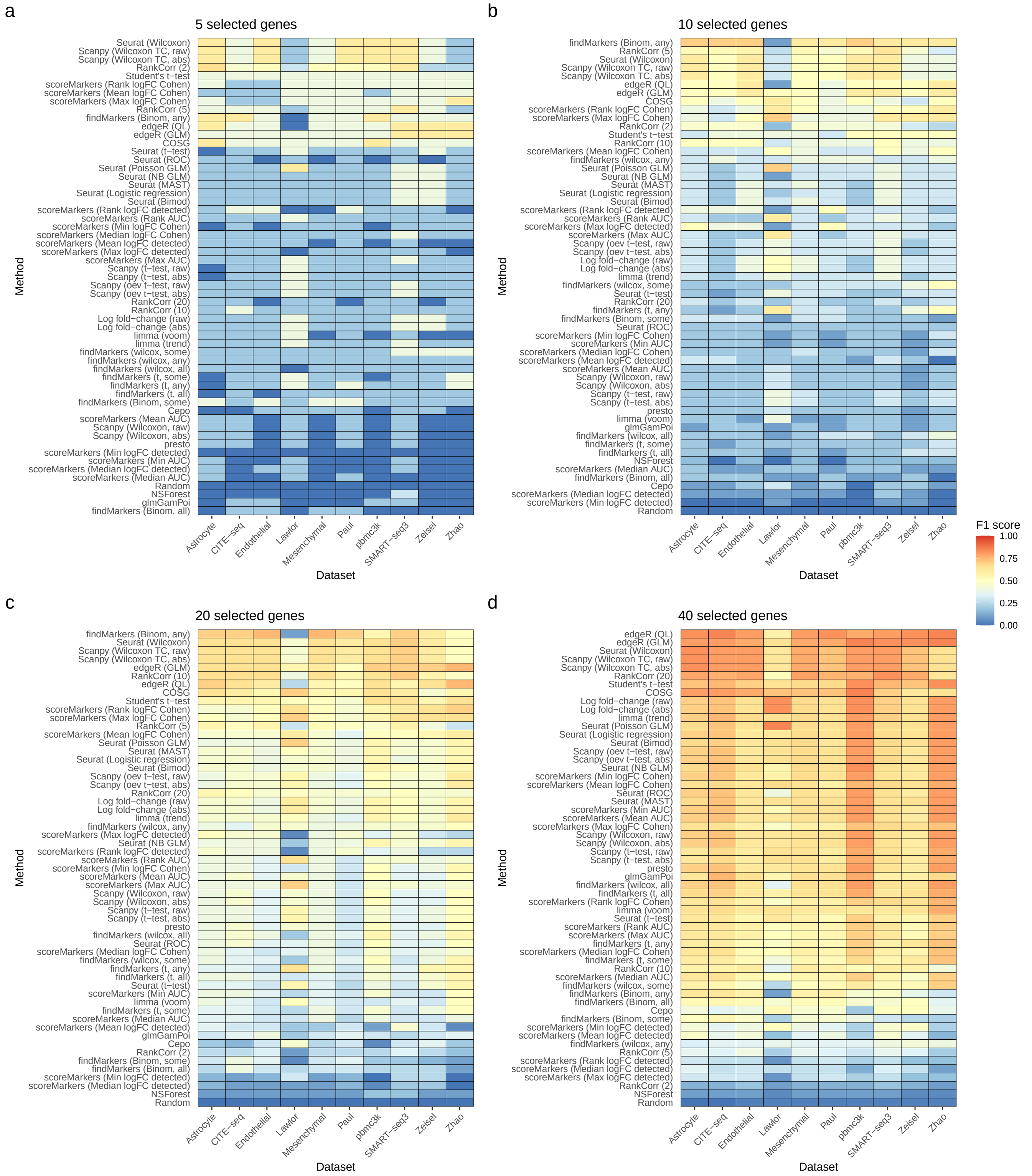


**Figure S10: F1 scores across simulations scenarios when the number of selected and true marker genes is varied** a) 5 genes are selected and used as true marker genes. b) 10 genes. c) 20 genes. d) 40 genes. When 40 genes are used the edgeR method shows increased performance.

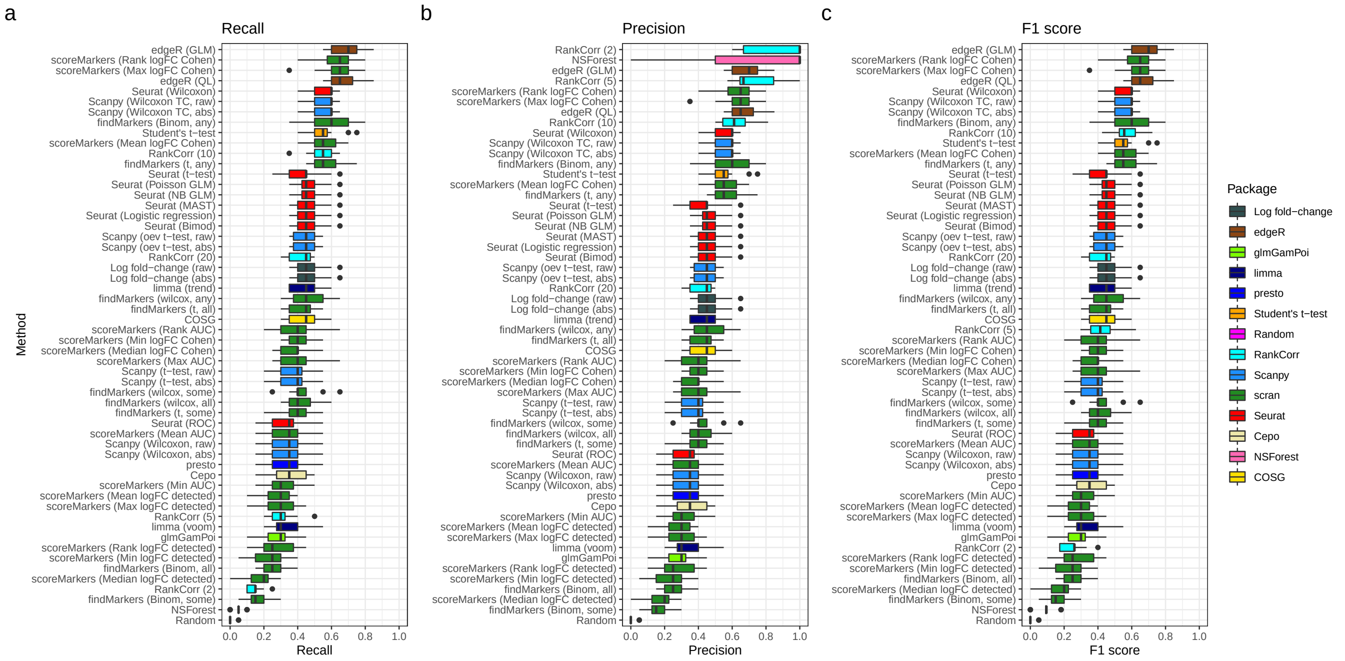


**Figure S11: Performance of all methods on simulated data in the Zeisel-based simulation scenarios** Boxplots show variations in performance across clusters and simulation replicates. a) Recall. b) Precision. c) F1 score.

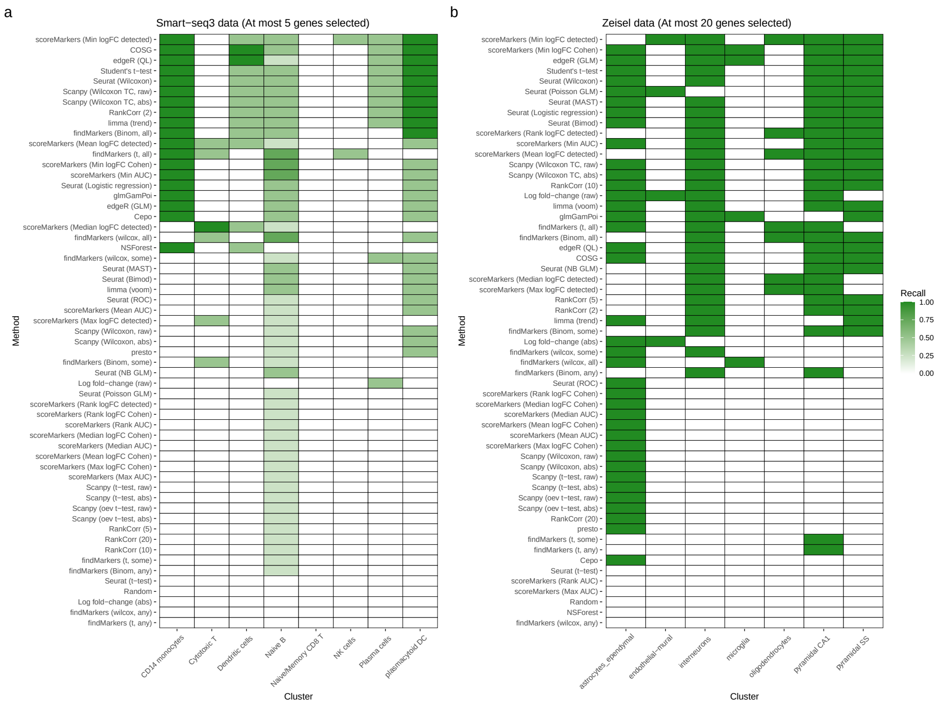


**Figure S12: Recall of methods on the Smart-seq3 and Zeisel datasets when recovering expert-annotated marker genes** a) Results for the Smart-seq3 dataset. At most 5 genes were selected and the expert-annotated marker genes were taken from the supplementary material of the paper describing the Smart-seq3 method. b) Results for the Zeisel datasets. At most 20 genes were selected and the expert-annotated marker genes were taken from the paper describing the Zeisel dataset.

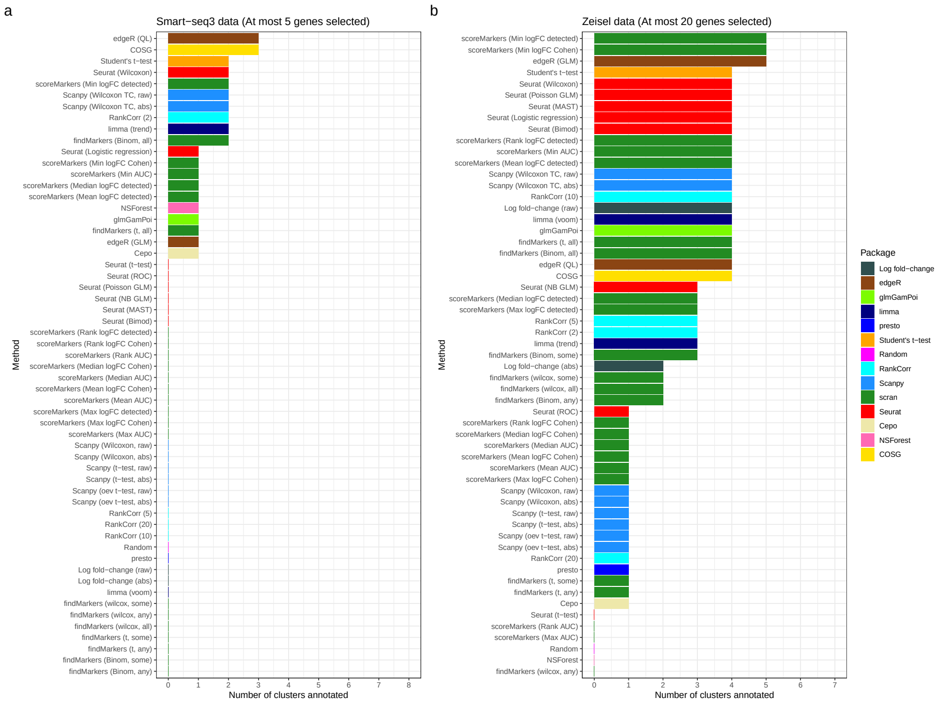


**Figure S13: Number of clusters annotated by methods for the Smart-seq3 and Zeisel datasets using expert-annotated marker genes** A cluster was taken to be successfully annotated if the method was able to recover all of its associated expert-annotated marker genes. a) Results for the Smart-seq3 dataset. At most 5 genes were selected and the expert-annotated marker genes were taken from the supplementary material of the paper describing the Smart-seq3 method. b) Results for the Zeisel datasets. At most 20 genes were selected and the expert-annotated marker genes were taken from the paper describing the Zeisel dataset.

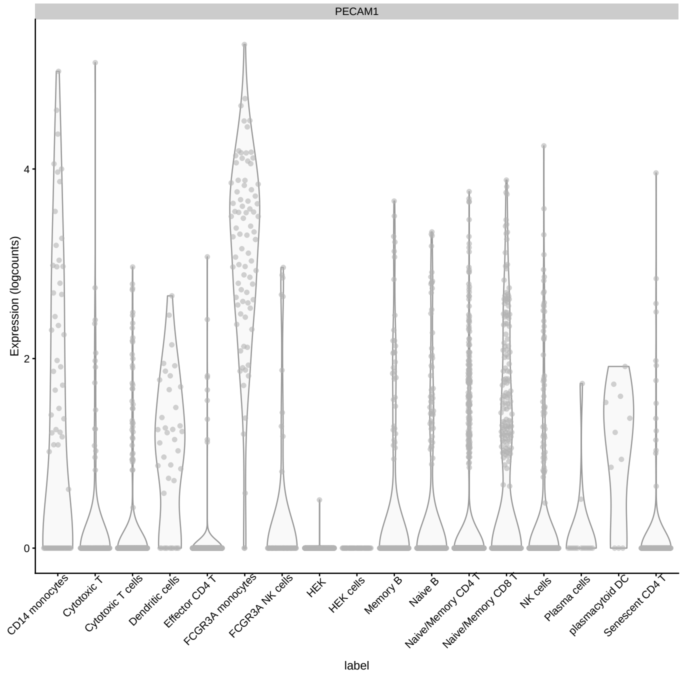


**Figure S14: Expression of the *PECAM1* gene across clusters in the Smart-seq3 dataset**

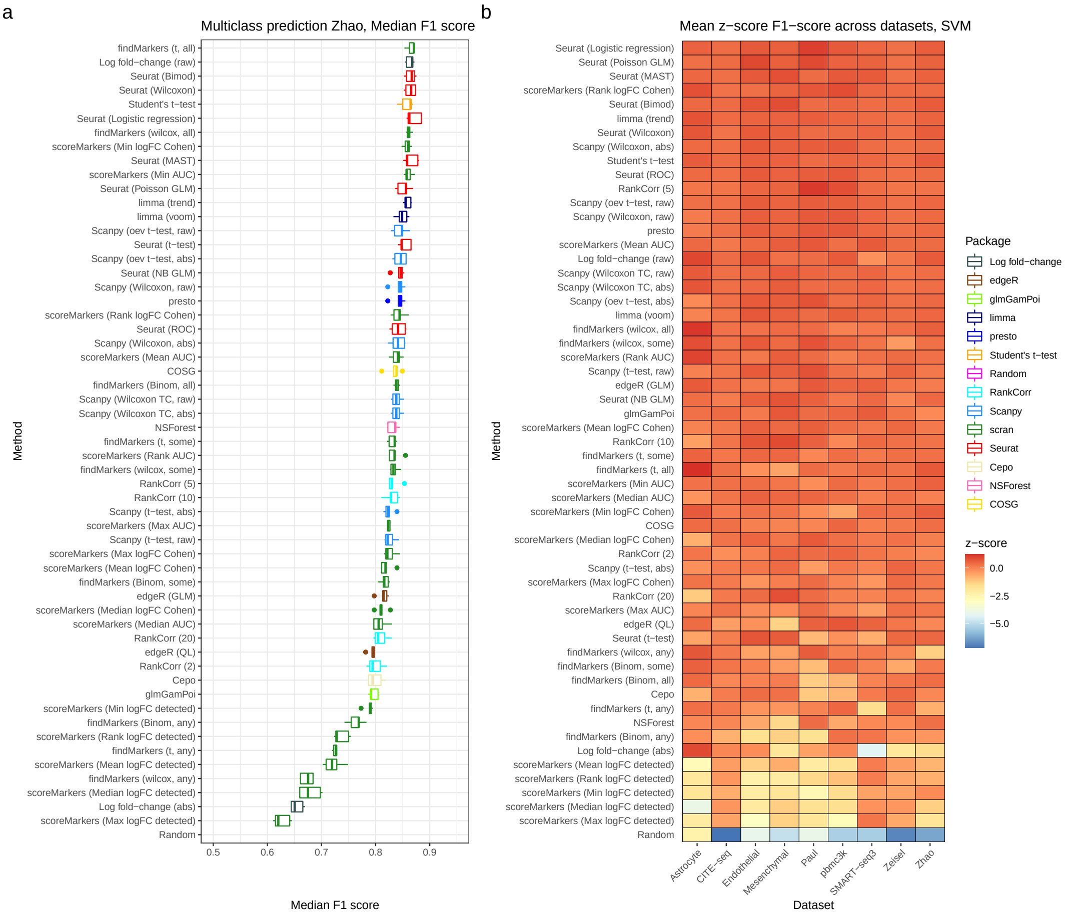


**Figure S15: Predictive performance measured using the SVM classifier** a) Median F1 score of the SVM classifer using genes selected by all marker gene selection methods in the Zhao dataset. Each point is the F1 score in one of the 5 folds. b) The z-score of the median F1 score of the SVM classifier (averaging across folds) in each dataset, for the classifier. Methods are ranked top to bottom by their mean z-score across datasets. Note that the Lawlor dataset is excluded from these results as the SVM classifier gave odd results when run on it.

**Table S3:** Expert marker genes: Zeisel

| Cell type | Marker gene |
|---|---|
| interneurons | PNOC |
| pyramidal SS | TBR1 |
| pyramidal CA1 | SPINK8 |
| oligodendrocytes | HAPLN2 |
| microglia | AIF1 |
| endothelial-mural | ACTA2 |
| astrocytes_ependymal | ALDOC |

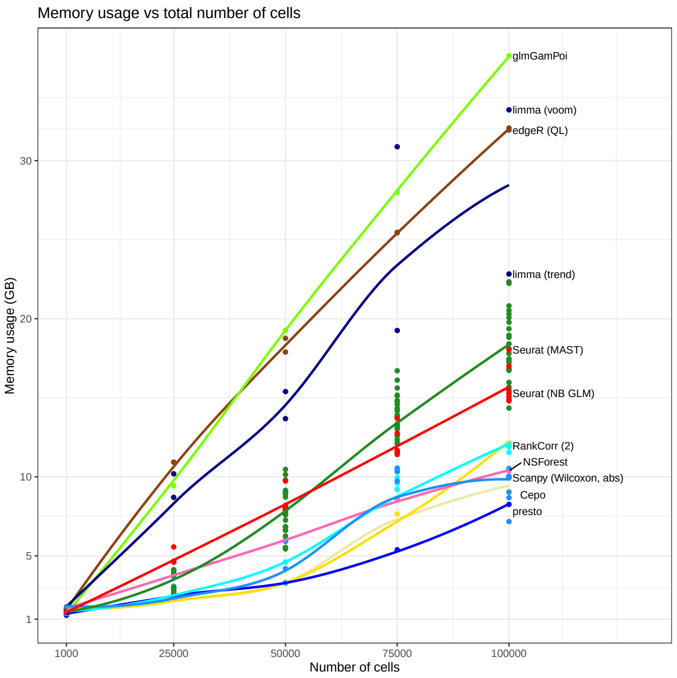


**Figure S16: Memory usage of methods on simulated datasets with varying number of total cells** All simulations parameters estimated from the pbmc3k dataset with DE factor location and scale parameters of 3 and 0.2 respectively. Datasets with 1,000, 25,000, 50,000, 75,000 and 100,000 total cells were simulated. All datasets had 5 clusters and 2000 genes.

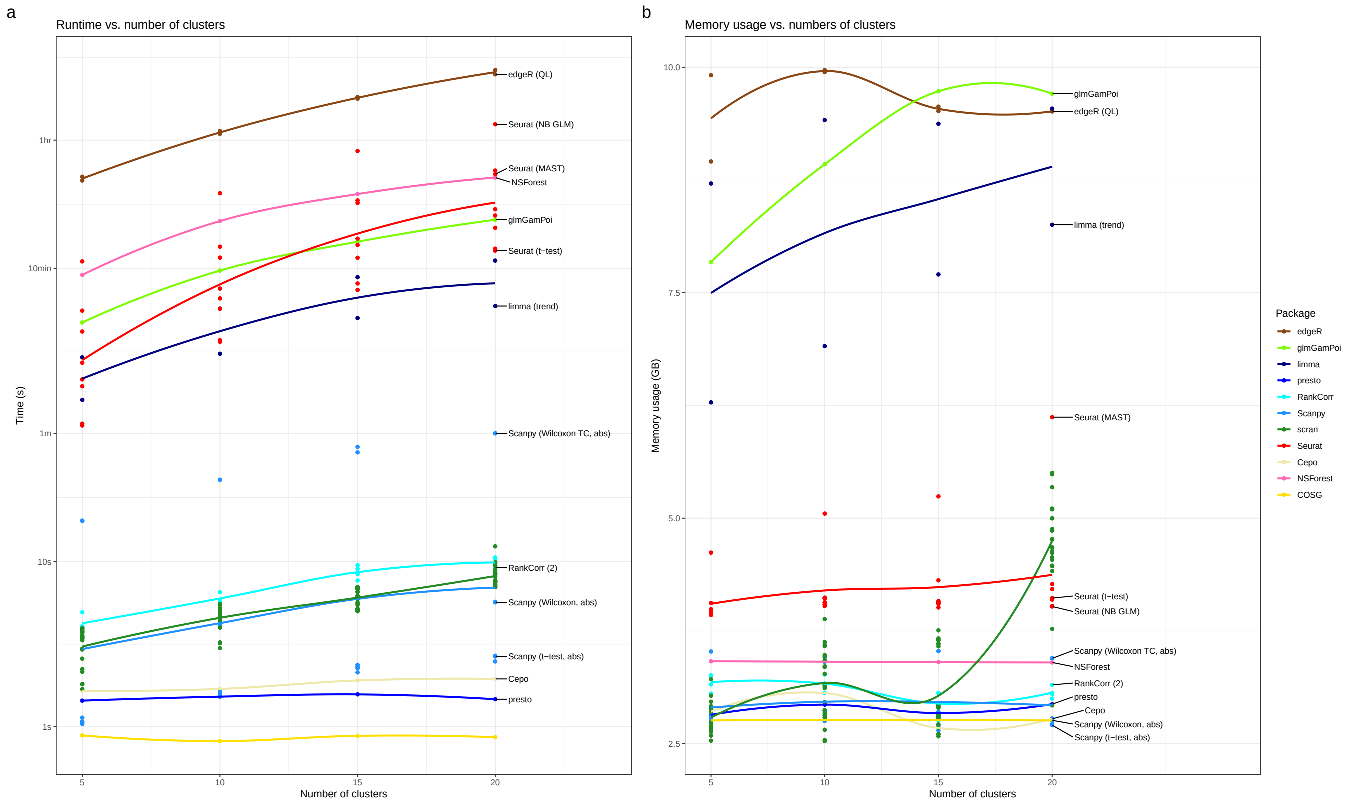


**Figure S17: Time and memory usage of methods on simulated datasets with different numbers of clusters.** All simulations had estimable parameters estimated from the pbmc3k dataset, DE factor location and scale parameters of 3 and 0.2, 20,000 cells and 2000 genes.

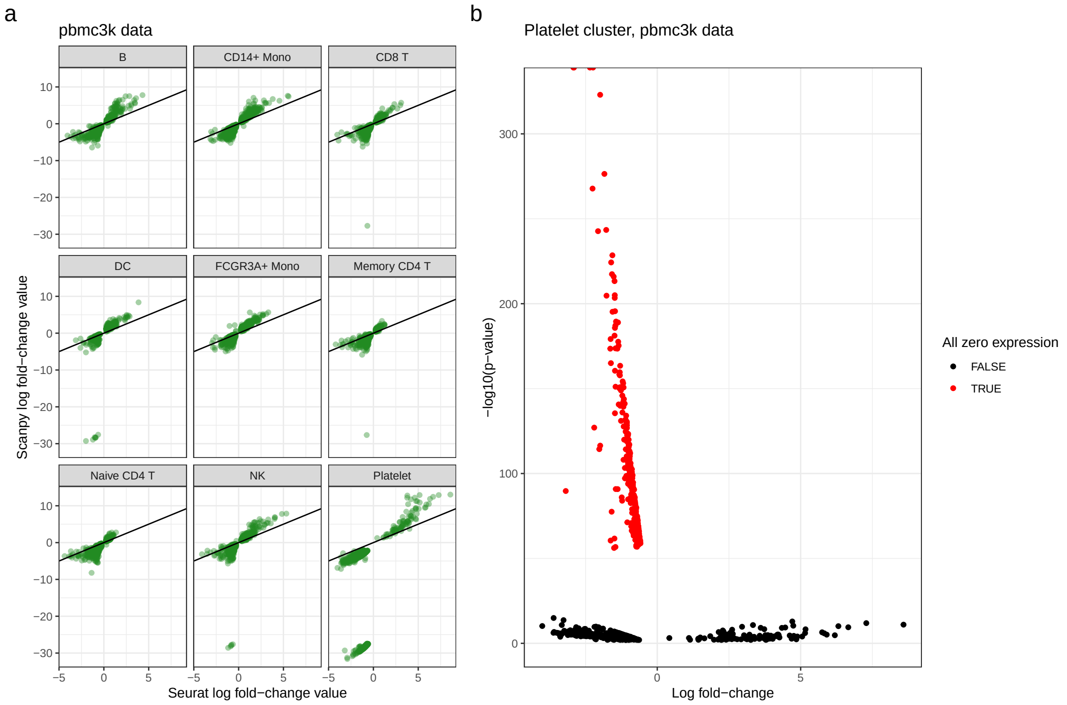


**Figure S18: Consequences of exactly zero expression** a) Scatter plots of Seurat and Scanpy calculated log fold-changes for each cluster in the pbmc3k dataset. Note the extreme values calculated by Scanpy for some clusters, such as the Platelet cluster. These occur when there is exactly zero expression a particular cluster. b) Volcano plot of genes for the Seurat t-test method in the Platelet cluster pbmc3k. A large number of genes with small log fold-change but highly significant p-values is caused by zero-expression in these genes in the Platelet cluster. Genes are coloured by whether they show zero expression in the platelet cluster.

**Table S4:** Expert marker genes: Smart-seq3

| Cell type | Marker gene |
|---|---|
| NK cells | NCAM1, KLRB1 |
| Naive/Memory CD8 T | PECAM1 |
| Cytotoxic T | GZMB, GZMA |
| Naive B | CD27, IGHM, IGHD, IL4R |
| Dendritic cells | KLF4, CD1C |
| CD14 monocytes | CD14 |
| plasmacytoid DC | IL3RA, TLR7 |
| Plasma cells | PRDM1, IRF4 |

**Table S5:** Dataset cluster filtering

| Dataset | Cluster |
|---|---|
| pbmc3k | B |
| Lawlor | Alpha |
| Zeisel | Oligodendrocytes |
| Paul | 14Mo |
| Smart-seq3 | Naive B |
| Endothelial | Endothelial - capillary |
| Astrocyte | Astro_HYPO |
| CITE-seq | 2 |
| Mesenchymal | Pericyte |
| Zhao | Naive B |
